# Distinct computations underlie movement error parsing during implicit motor adaptation and perturbation detection

**DOI:** 10.64898/2026.06.25.734533

**Authors:** Hyosub E. Kim, Jack O. Darley, Michael S. Landy, Romeo Chua, Dusty J. Fox

## Abstract

The human sensorimotor system is remarkably effective at automatically parsing total movement error into its constituent parts, the error component due to a perturbation, or externally-generated error (EGE), versus the error component due to motor noise, or internally-generated error (IGE). This implicit error parsing, and its associated perceptual processes, directly contrasts previous work showing that humans will explicitly misattribute motor errors caused by relatively large external perturbations to their own motor variability. While the combined results suggest a dissociation between perception for action—which allows for precise and automatic error parsing—and perception for conscious detection, this must be inferred across studies using different methodologies. Here, we combined a within-subjects study design and computational modeling to shed light on the principles underlying implicit adaptation to a perturbation and explicit perturbation detection. Neuro-typical adults participated in two experiments consisting of pseudo-randomized rotations during reaches to a single target, with one session requiring explicit reports after each reach of whether a perturbation was detected. Participants demonstrated a clear dissociation between implicit responses to a perturbation and explicit detection, with robust adaptation to 1° EGEs, but an inability to reliably report the presence of an EGE until it reached ∼ 4°. Our modeling results explain this behavioral dissociation as a consequence of the sensorimotor system applying distinct computational strategies to these behaviors. For the adaptation task, the best-fitting model assumes the participant compares proprioceptive and visual cues to detect a perturbation and corrects for a proportion of this error. For signal-detection, the best-fitting model used Bayesian causal inference in which sensory cues are optimally integrated with a prior on their cause. Combined, our study sheds light on the dichotomy between perception for action versus perception for recognition in the context of visuomotor control and adaptation.

**Author Summary:** Humans are remarkably skillful at adapting voluntary movements to novel contexts and environments. Much of this motor flexibility depends on accurately determining whether a movement error was caused by an external perturbation or by intrinsic motor variability, the former requiring adaptation while the latter should be ignored. Here, we investigated the relationship between implicit motor adaptation to visual perturbations (rotated feedback of reaching movements) and our ability to consciously detect those same perturbations. Through a combination of behavioral testing and mathematical modeling, we show a striking dissociation between the two behaviors. Participants implicitly adapted their reaches in response to visual rotations as small as 1° but were unable to reliably report the presence of a rotation until it was nearly four times as large. Our modeling work indicates that two distinct strategies underlie implicit motor adaptation and explicit perturbation detection. For adaptation, small discrepancies between vision and proprioception, our sense of where our body and limbs are in space, automatically trigger an adaptive response. In contrast, for explicit perturbation detection, participants combined noisy sensory information with prior expectations about motor performance and the environment in a statistically optimal manner.

## Introduction

Imagine a hurried weekday morning and quickly reaching for, and missing, the power button on your coffee machine. Holding a smartphone while reaching may have subtly changed your normal reaching dynamics and caused this movement error, in which case your motor system should automatically adapt its sensorimotor mapping to account for this systematic “perturbation” on the next try. On the other hand, if your miss was simply due to intrinsic motor variability, the motor system should avoid adapting future movement in response to this random error since continuing to learn from noise leads to behavioral instability and an amplification of future errors (Faisal et al., 2008; van Beers, 2009). Given the ecological importance of discriminating between different types of movement error, perhaps it is not surprising then that humans are remarkably accurate and precise in automatically parsing total movement error into its constituent parts: the error component due to a perturbation, or externally-generated error (EGE), versus the error component due to motor noise, or internally-generated error (IGE) (Fig 1b). Humans demonstrate robust implicit adaptation to very small (2°) EGEs in the form of randomized visuomotor rotations while ignoring identically-sized errors caused by motor noise (IGE), operationally defined here as the deviation between the hand and the aim location (i.e., the target) (Kim et al., 2025; Ranjan & Smith, 2018). This effective movement error parsing is thought to be due to the sensorimotor system’s ability to accurately predict hand trajectory, based on an efference copy of the motor command (Carriot et al., 2013; Ranjan & Smith, 2018; Sommer & Wurtz, 2008; Szarka et al., 2025). Our recent work extended this idea by formalizing how a participant should optimally combine sensory cues, including motor prediction-based contributions to proprioception, to infer the sources of movement error (Kim et al., 2025).

**Figure 1:**
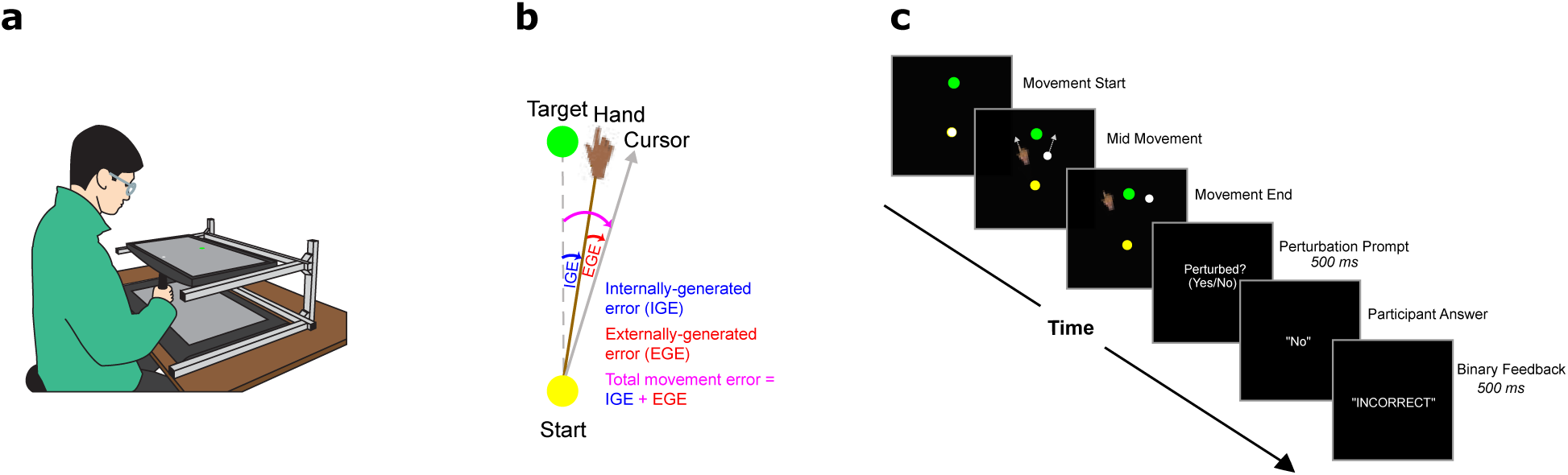
(a) Experimental set-up. Participants made quick point-to-point reaches by sliding a stylus along a graphics tablet. Vision of the hand was absent due to being in a darkened room and the horizontally-oriented monitor above the tablet. (b) Decomposition of total error into externally-generated error (EGE) and internally-generated error (IGE) components. IGE is equivalent to motor noise, or how far off the intended aim (i.e., the target) the reach was. EGE refers to the external perturbation—in this case, the magnitude of the visuomotor rotation. Adaptation is quantified as the difference between reach angles on post-versus pre-perturbation trials. (c) Signal-Detection Task trial sequence. The target is represented by a green circle and the cursor by a white circle. The hand is shown in the diagram for illustration purposes only. Participants indicated with a mouse click controlled by their non-reaching hand whether they thought visual feedback was perturbed or not and received binary feedback regarding their choice.

Returning to our example, what relationship does awareness of the changed dynamics caused by the smartphone have to implicit adaptation? This impressive capacity to accurately discriminate between the smallest contributions of IGE and EGE to movement error, and *implicitly* adapt, is in direct contrast to our seemingly poorer ability to *explicitly* detect the presence of an EGE. Humans need to experience visuomotor rotations that are at least 1.5*x* the standard deviation of their motor variability, roughly 4° or more, to reliably report their presence (Gaffin-Cahn et al., 2019). The combined results across these studies suggest a potential dissociation between perception for action—which underlies such precise and automatic error parsing during implicit adaptation—and perception for conscious detection, as implicit adaptation to EGE occurs for errors that fall well below the threshold for explicit detection. However, this potential dissociation must be inferred across different studies using substantively different methodologies. More directly, it is unknown whether such a high explicit detection threshold exists when conditions across tasks are more closely matched. It remains an open question whether a higher explicit detection threshold is due to task-dependent changes in the accessibility of sensory information to explicit reporting, or a qualitatively different computation during implicit adaptation versus explicit detection, similar to the sharp distinction between implicit adaptation and explicit strategies (e.g., re-aiming) employed during sensorimotor learning tasks (Bond & Taylor, 2015; Kim et al., 2021; Miyamoto et al., 2020; Tsay et al., 2024).

Here, we combined computational modeling and a within-subjects study design to shed light on the computational principles underlying implicit adaptation to, and explicit detection of, movement perturbations. Neuro-typical adults participated in two separate experimental sessions, with the order counter-balanced. We designed the study to have two experiments in order to optimize participant performance on each individual task (see Methods). During the adaptation task, participants experienced pseudo-randomized rotations while making fast point-to-point reaches to a single target. During the signal-detection task, participants experienced the same range of perturbations as during the adaptation task, but this time were instructed to explicitly report after each reach whether they detected the presence of a perturbation or not. To formalize our understanding of these behaviors, we designed three distinct computational models that we fit to our data. The Parsing of Internal and External Causes of Error (PIECE) model is a Bayesian causal-inference model in which the observer optimally combines visual, proprioceptive and, presumably, motor-predictive cues to compute a posterior regarding whether sensory feedback was perturbed or not (Kim et al., 2025). In this model, adaptive motor output reflects an average of rotation estimates in the absence and in the presence of a perturbation, weighted by the inferred posterior probabilities of each scenario. (N.B.: Despite modeling perceptual and motor behaviors, we continue to use the term ‘observer’ in reference to our theoretical decision-makers, as opposed to ‘actor’ or ‘agent’, to maintain consistency with the terminology associated with the perceptual models we utilize in this study.) The other two models were adapted from a previous explicit-detection study (Gaffin-Cahn et al., 2019). The Comparison Observer (CO) model compares the discrepancy between visual and proprioceptive cues to decide whether there was a perturbation or not, whereas the Visual-Cue-Only (VCO) model assumes the participant utilizes only vision to make their judgment.

We report a clear behavioral dissociation between implicit adaptation and explicit detection: Participants showed robust implicit adaptation to EGEs as small as 1° and no adaptation to IGEs, yet required a perturbation of ∼ 4° before reliably reporting its presence. Adaptation was best captured by the CO model, supporting the idea that the computational goal of the adaptation system is to estimate and correct for any external perturbation. In contrast, the PIECE model best explained behavior on the signal-detection task, indicating participants optimally combined priors with incoming sensory information when having to make a binary decision regarding the presence of a perturbation. Combined, our results demonstrate that implicit adaptation and explicit perturbation detection are two dissociable behaviors governed by computationally distinct strategies.

## Results

### Differential Adaptation To IGE-EGE

To determine the exact nature of how implicit error parsing is related to explicit perturbation detection, we conducted a within-subjects design study involving 16 healthy, neurotypical young adults who were tested on a visuomotor-adaptation task and a perturbation-detection task. While the order of experiments was counter-balanced across participants, we first describe the results of the adaptation task.

Participants made fast point-to-point reaches to a single target while controlling a small white cursor. After 100 baseline reaches (to provide an estimate of movement variability), depending on the trial, the cursor could be rotated by 0°*, ±*1°*, ±*2°*, ±*3°*, ±*6°, or *±*10° (70 trials per perturbation level for a total of 770 trials). While previous studies used only *±*2° and *±*4° rotations, we added the extra perturbation sizes in order to ensure we could assess both implicit adaptation and perturbation detection with perturbations that were below and above the explicit perturbation-detection thresholds. This range of perturbations also roughly matched those used by Gaffin-Cahn and colleagues (2019). Each rotation trial was always surrounded by null (no rotation) trials, with the rotation magnitude pseudo-randomized across the experimental block (Fig 1 shows the experimental set-up). Consistent with prior work (Kim et al., 2025; Ranjan & Smith, 2018), half of the null trials (randomly selected) had no visual feedback, which increased our sensitivity to detect adaptation to IGE, if present (see Methods). The no-visual-feedback trials were treated identically to the null trials with visual feedback in our analyses, since the presence or absence of visual feedback should have no influence on reach angle at maximum radial velocity (our primary dependent variable). Additionally, in our analyses, the overall motor error was decomposed into its internally and externally generated parts. The internally-generated error (IGE) is defined as the random error that occurs on every reach (i.e., any angular deviation of the hand trajectory from the target), and on unperturbed reaches, serves as the only contribution to total error. IGE is the result of intrinsic motor variability plus any intrinsic bias in reach direction. The latter is the residual error in reaching that participants appear to tolerate, thought to arise from errors in sensory coordinate transformations (Wang et al., 2026).

Externally-generated errors (EGE) are caused by external perturbations (e.g., visuomotor rotations) and are under experimental control (Fig 1). In this experiment, EGEs were defined as the visuomotor rotations. With each perturbation trial being surrounded by null trials, adaptation was quantified as the difference in reach angles on trials immediately following versus immediately preceding a perturbation trial: adaptation*_t_* = reach*_t_*_+1_ *−* reach*_t−_*_1_, where *t* indexes the perturbation trial. This allowed us to estimate the adaptation from one perturbation trial while discounting any carryover from the previous perturbation trial. Combined, these methods allowed us to easily dissociate the influence of IGE versus EGE on implicit adaptation on a trial-by-trial basis (Kim et al., 2025; Ranjan & Smith, 2018).

Consistent with prior reports, all participants showed highly accurate error parsing, robustly adapting to all non-zero rotations, while discounting size-matched IGEs. A representative participant’s data are shown in Fig 2. (Figures of all individual participant data are provided in the Supporting Information S1 and S2 Figs.) As seen in Fig 2, there also appeared to be a saturation of error corrections for rotations of 6° or larger, consistent with prior reports (Kim et al., 2018; Morehead et al., 2017). To provide evidence for this saturation, we compared simple linear regression slopes using all of the data (-0.32 [-0.35, -0.29]; mean [95% bootstrapped CI]) as compared to when leaving out the *±*10° rotation data (-0.40 [-0.44, -0.35]) and found reliably lower slopes when all data were included (*t*_15_ = 7.56, *p* = 1.71 *×* 10*^−^*^6^). A number of proposed explanations of saturated adaptation have been put forward, which we elaborate on in the Discussion. Regardless of its source, this behavior serves as another indication that adaptation during the task is implicit, as explicit strategies are known to scale linearly for all perturbation sizes (Bond & Taylor, 2015; Hutter & Taylor, 2018).

**Figure 2:**
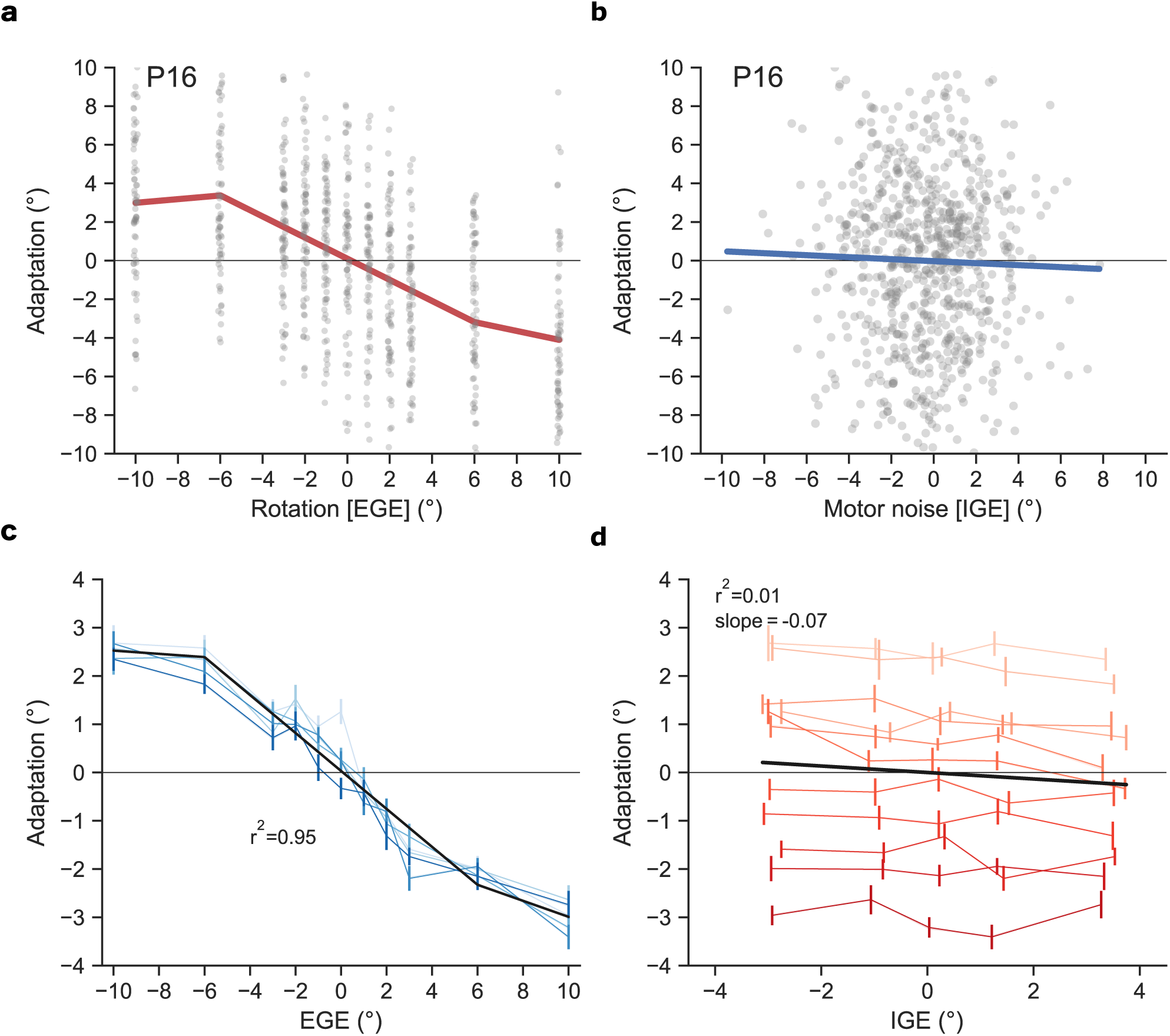
(a) Data for a representative participant (Participant 16) showing adaptation as a function of EGE (gray dots are single-trial adaptive responses). (b) The same adaptive responses plotted as a function of IGE. (c-d) Population-averaged adaptive responses were binned based on the level of IGE (shade of blue; lightest shade represents most negative IGE, i.e., 1st quintile, darkest represents most positive IGE, i.e., 5th quintile) and EGE (shade of red; lightest shade represents *−*10° EGE and darkest shade represents +10° EGE). The error bars in (d) are centered on x-axis locations corresponding to the mean bin center across participants. The black lines represent the mean adaptive response as a function of EGE in (c), and as a function of IGE in (d). The vast majority of the variance in adaptive responses (95.1%) was explained by EGE, while IGE accounted for only 1% of the variance. Error bars are 95% bootstrapped confidence intervals.

To analyze data at the group level, we performed a binned analysis. Data were first binned based on EGE size, and then for each of the 11 EGE sizes (including 0°). We further binned the data based on the level of IGE by splitting IGEs into quintiles (see Methods), resulting in 55 total bins. This procedure was applied separately for each participant. The results of these analyses are presented in Fig 2c-d, where shades of blue represent IGE quintile and shades of red represent EGE size. In this figure, where the blue functions represent the mean adaptive response (and 95% CI) across participants for each IGE quintile at every perturbation level, and the black line represents the best-fit piecewise linear-regression across these data. This figure shows that EGE was almost completely predictive of the adaptive response, as the different levels of IGE had little influence on adaptation magnitude. This was further supported by the very high amount of variance explained, 95%, even after adjusting for the 5 free model parameters (3 slope parameters, 1 bias term, and 1 parameter used to set the break point at *±*6°). We note that when fitting these same data with a linear function, the variance explained was still above 92%. In stark contrast to Fig 2c, in Fig 2d we see that IGE had no relationship to adaptation. These data were clearly split by EGE magnitude, represented by the red functions, and across each IGE quintile, each red function is relatively flat. There was no systematic relationship between adaptation and IGE, and only a negligible amount of the variance in the EGE/IGE grid can be explained by IGE (black line, *r*^2^=0.01).

These results are another demonstration of how humans are able to discriminate between EGE and IGE and are made more impressive by the robust adaptation observed with EGEs as small as 1° (Fig 2a,c). To quantify implicit adaptation to the 1° rotations, we performed a bivariate regression on each participant’s data using adaptation measures from *−*1°, 0°, and + 1° trials, using EGE and IGE as predictor variables. There was a robust response to these tiny perturbations (EGEs), with highly reliable slopes, i.e., sensitivities (*β*_EGE_ : *−*0.50[*−*0.60*, −*0.40]; *t*_15_ = *−*9.65, *p* = 7.98 *×* 10*^−^*^8^). In stark contrast, the sensitivities to IGE in this subset of data hovered close to zero (*β*_IGE_ : *−*0.07[*−*0.13*, −*0.01]; *t*_15_ = *−*2.18, *p* = 0.05).

### Perturbation-Detection Thresholds

To assess participants’ explicit perturbation-detection thresholds, the same 16 participants experienced identical rotation sizes as during the adaptation task, but were now instructed to explicitly report after each reach (800 total; full randomization of 400 trials with perturbation—40 per non-zero perturbation level—and 400 without perturbation) whether they detected the presence of a perturbation or not. Briefly, immediately following completion of their reach, participants saw a screen asking them if they thought they were perturbed or not. They indicated their responses with a mouse-button press and subsequently received binary feedback regarding the accuracy of their judgment. To quantify explicit detection thresholds, we first calculated *d’* (“d prime”) values for each participant and each perturbation magnitude, combining data from negative and positive-signed rotations. The value of *d’* provides a standardized metric for discriminability between signal and noise, i.e., the signal-to-noise ratio. As the relationship between *d’* and perturbation magnitude appeared linear, we fit each individual’s data with a linear regression model. We then utilized the linear fits to interpolate the detection threshold, which we define as the perturbation magnitude resulting in a *d’* value of 1, translating to approximately 69% correct responses for a neutral criterion.

In comparison to the adaptation results, these same participants demonstrated greater difficulty explicitly reporting the presence of a perturbation. We observed detection thresholds that were ∼ 1.5*x* baseline motor variability, quantified as the SD of the last 50 baseline trials (predicted *d^′^* = 1 for a rotation of 3.62° [3.36°, 3.94°]; *σ*_motor_ : 2.07° [1.86°, 2.26°]); Fig 3), consistent with the results of Gaffin-Cahn and colleagues (2019). This consistent relationship across studies suggests that, in terms of absolute units, participants with greater motor output variability may be more willing to attribute larger visual errors to themselves than to an external source. While there was a positive relationship between the standard deviation of baseline reaching and detection threshold, this correlation was not reliable (*r*^2^ = 0.16; *p* = 0.13). Regardless, the combination of robust implicit adaptation to EGEs as small as 1° with explicit detection thresholds of almost 4° supports the idea that perception for action and perception for conscious detection are distinct processes.

**Figure 3:**
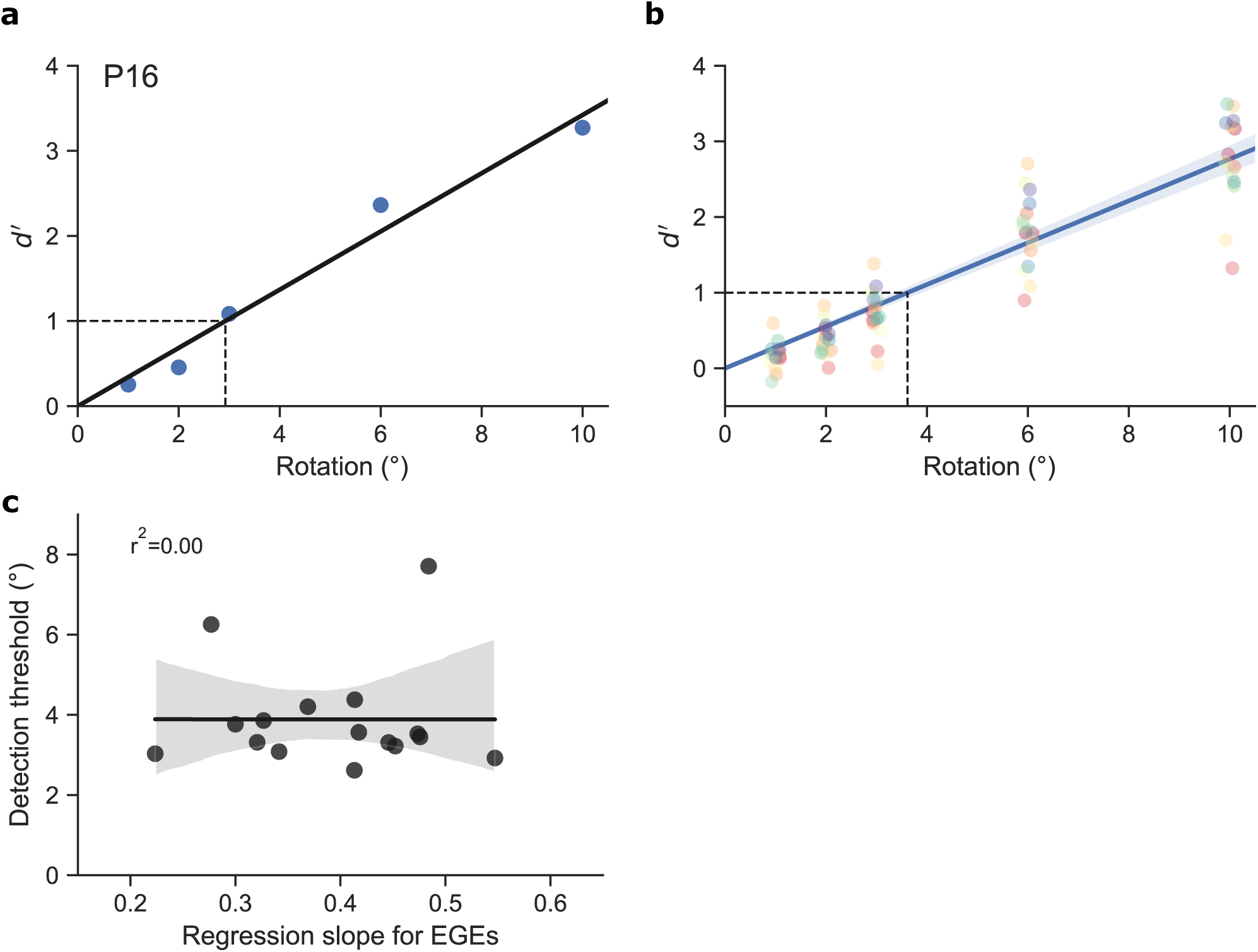
(a) Data for a representative participant (Participant 16) from the signal-detection task. Discriminability (*d^′^*) increased linearly with perturbation size. (b) Group-level data for the signal-detection task. On average, *d^′^* = 1 (69% hit-rate for a neutral criterion) when the perturbation is 3.62°. (c) Correlational analysis between error sensitivity measures during the adaptation and signal-detection tasks. Note that the regression slopes have been sign-flipped for ease of interpretation (higher value indicates higher sensitivity to EGE). Dots represent individuals. Shaded error represents the 95% confidence interval.

To more rigorously examine whether there was a relationship between an individual’s sensitivity to EGE during the adaptation versus signal-detection tasks, we conducted a correlation analysis. We quantified implicit sensitivity to EGE with the *β* values of adaptation to EGE from best-fit lines to the -6°:+6° data, while explicit sensitivity was quantified using interpolated perturbation values corresponding to detection performance of *d^′^* = 1. If sensitivities across tasks were related, we would expect to see strong anti-correlation, as that would indicate those individuals with higher sensitivity to EGE during adaptation would show a lower detection threshold during the signal-detection task. However, we observed no relationship between sensitivity to EGEs during adaptation and estimated thresholds for the perturbation magnitude corresponding to a *d’* of 1 (*r*^2^ = 0.00; *p* = 1.00).

As further confirmation of the dissociation between implicit adaptation and explicit perturbation detection, we also examined the exact same relationship using data from only the signal-detection task. This is not as robust an analysis as the comparison across tasks, due to the experiment being optimized for signal detection as opposed to implicit adaptation (see Methods). Despite these caveats, we ran the correlational analysis and again found no relationship between sensitivity to EGEs and perturbation-detection thresholds within the same task (*r*^2^ = 0.03; *p* = 0.51).

Combined, these results indicate a decoupling between implicit adaptation and explicit error detection, with the adaptation system being highly sensitive to EGEs falling well below explicit-detection threshold.

### Model-Based Analyses

To formalize our understanding of these behaviors, we designed three distinct computational models with which to fit our data. We also took a simulation-based approach to characterizing the alignment between model predictions and actual behavior through posterior-predictive checks (R. C. Wilson & Collins, 2019). Specifically, after finding the maximum-likelihood estimates (MLEs) of model parameters for each individual and task, we then ran simulations of each model through the exact behavioral protocols using those estimated parameter values (as in Fig 5).

The Parsing of Internal and External Causes of Error (PIECE) model, which successfully explained error parsing during implicit adaptation in our previous work (Kim et al., 2025), is similar to the ideal observer model used by Gaffin-Cahn and colleagues (2019). Notably, we have formalized the model such that it readily accounts for implicit adaptation and explicit perturbation detection. PIECE frames error detection (and adaptation) as a process of causal inference whereby the observer optimally integrates noisy visual and proprioceptive cues to determine the probability of a perturbation. It performs inference based on the generative model of perturbation shown in Fig 4, where *C* is the causal node which gives rise to perturbed and unperturbed world states. The observer utilizes visual and proprioceptive cues (*x_v_* and *x_p_*) dependent on the hand position (*x*_hand_) during reaching. We assume proprioception is aided by motor prediction, but we are unable to disambiguate the two. This is described in Methods, where a full derivation of the model can be found. In stark contrast, the Visual-Cue-Only (VCO) model posits that errors are detected only when the noisy visual cue regarding hand position, which includes the effect of a rotation if present on that reach, deviates sufficiently from the participant’s own intrinsic motor bias (if zero motor bias, this would equal the sensed distance of the cursor from the target). The motor bias is the mean signed error, on the order of 1 *−* 2° that the sensorimotor system is known to tolerate, with potential origins in a misalignment between proprioceptive and visual coordinate frames (Wang et al., 2026). Even during baseline reaching, when there is no perturbation, participants do not fully resolve this error, and so the bias is treated as a free parameter in all of our model fits. The Comparison Observer (CO) model detects an error only when the difference between the visual and proprioceptive cues exceed a participant-specific threshold (*D*_threshold_). The VCO and CO models were originally designed for explicit error detection only; thus, to fit adaptation data, we incorporated ideas from standard state-space models of adaptation (Smith et al., 2006; Thoroughman & Shadmehr, 2000), modifying both models such that adaptation occurs specifically in response to detected errors, with the magnitude of adaptation being a fixed fraction of this error. As in the standard state-space model, the trial-by-trial error correction is *βe_t_*, where *β* represents the proportion of the error, *e_t_*, that is corrected for. Here, rather than defining *e* as the sensory-prediction error (the difference between actual and predicted visual feedback), *e* is determined by the respective model definitions of error (see Methods for full equations). In contrast to VCO and CO models, fitting PIECE to both perceptual and motor tasks is feasible without any model modifications, one of the inherent strengths of the Bayesian causal-inference framework.

**Figure 4:**
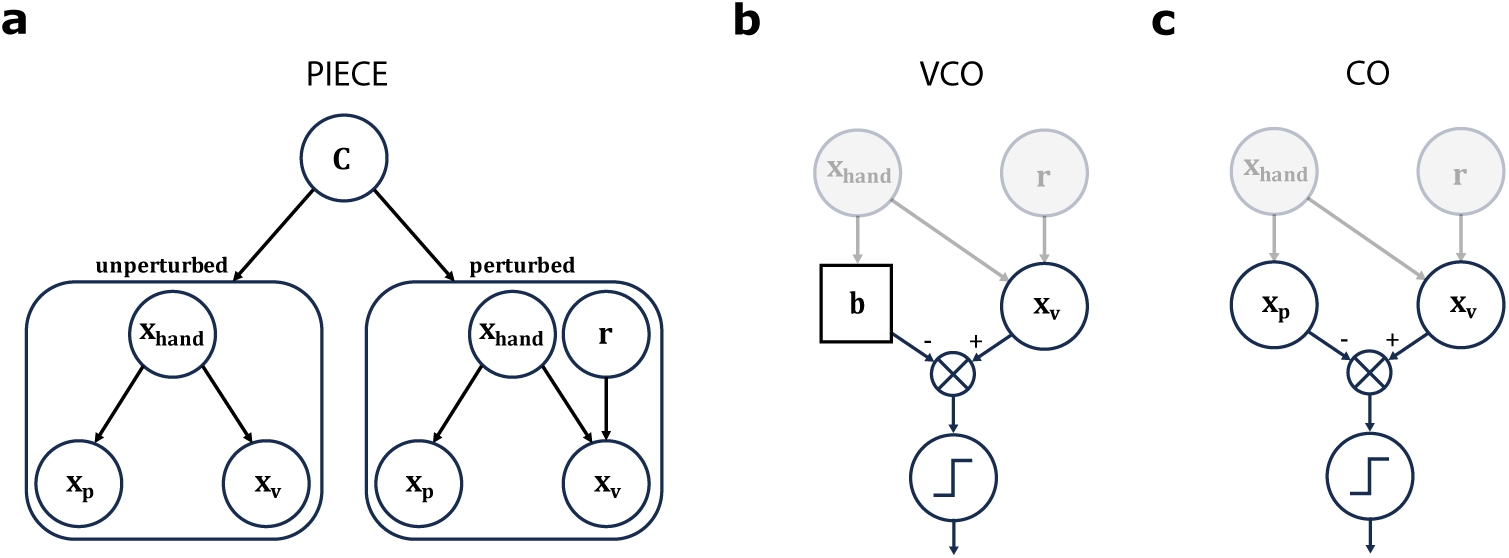
Graphical representations of computational models. (a) Graphical generative model for PIECE. During adaptation, PIECE proposes the participant computes a posterior on Cause, *C*, based on visual and proprioceptive cues, to weight their estimate of the rotation magnitude, *r*. During signal-detection, inference of *C* forms the basis for their psychophysical judgments, with detection occurring when the posterior probability of perturbation is greater than or equal to 50%. (b) Schematic for VCO. In the VCO model, the observer compares the distance between the visual cue and their own intrinsic motor bias, a model parameter (c) Schematic for CO. In the CO model, the observer compares the distance between visual and proprioceptive cues. For (b) and (c), if the distance exceeds an internal threshold, the observer detects the perturbation (represented by step function in bottom node) and adapts their subsequent reach. Note that for both models, the nodes for *x*_hand_ and *r* are shaded gray to indicate that, in contrast to PIECE, the observer does not perform inference on these variables.

We fit each individual’s adaptation and signal-detection datasets using maximum-likelihood estimation, and computed Akaike Information Criterion (AIC) scores for objective model comparison. The first research question we attempted to answer through our model fitting procedures was whether implicit adaptation and explicit perturbation detection share the same underlying computation. This is answered by model selection comparing joint fits with a shared set of MLEs versus separate fits to each task. Therefore, we first compared the quality of fits to each participant’s data when optimizing only one set of free parameters per model per participant versus optimizing two sets of parameters per model (one for each task). The joint fits to both tasks favored PIECE over VCO for 13/16 participants, and for 12/16 participants against the CO model. While this suggests PIECE is the most well-supported model if the goal is to find a single modeling framework to explain both tasks, AIC scores highly favored separate fits for each task in 15 out of the 16 total participants. This result allows us to confidently rule out the possibility that implicit adaptation and explicit perturbation detection share the same computations. This is not surprising given the behavioral dissociation observed across the two tasks.

Our second model-based question was whether adaptation and detection at least share the same computational strategies. That is, does one model (with two separate parameter sets) best fit data from both tasks, or do participants utilize different computational strategies depending on the task? This question is best answered by model selection comparing fitting all nine possible combinations of model pairs across both tasks (i.e., PIECE:PIECE; PIECE:VCO; PIECE:CO; VCO:PIECE, VCO:VCO; etc.) Based on the inferior model performance from the joint fit and our research questions, we focus our reporting here on the model-comparison results from separately fitting each task.

The CO model unanimously outperformed both the PIECE and VCO models on the adaptation task. PIECE was second best, being judged better than VCO on adaptation for 12/16 participants. We initially expected PIECE to best fit the adaptation data, given its superior performance in our previous study against several other computational models of adaptation from the literature (Kim et al., 2025). Despite these results, we note here that CO and PIECE share several conceptual similarities, such as optimally combining sensory information to infer the rotation magnitude. In fact, the CO model can be viewed as a heuristic-like version of PIECE, as it does not compute a posterior or concern itself with full probability distributions. Both of these models are in stark contrast to the VCO model, which posits that participants only use visual information to correct for motor errors. Data from a representative participant are shown alongside simulations using that participant’s best-fit parameters (i.e., posterior predictive checks). Fig 5a shows that CO and PIECE both capture the salient behavioral features in the adaptation task of effective error parsing and a linear response for small perturbations. Although the CO model had greater difficulty accommodating saturated adaptive responses for 6 and 10° perturbations for reasons we discuss later, this did not prevent it from being judged superior to PIECE. The combination of decisive support for CO over PIECE based on information criteria, combined with lack of a clear, qualitative difference between CO’s and PIECE’s posterior predictions is something we discuss in greater detail in the Discussion.

**Figure 5:**
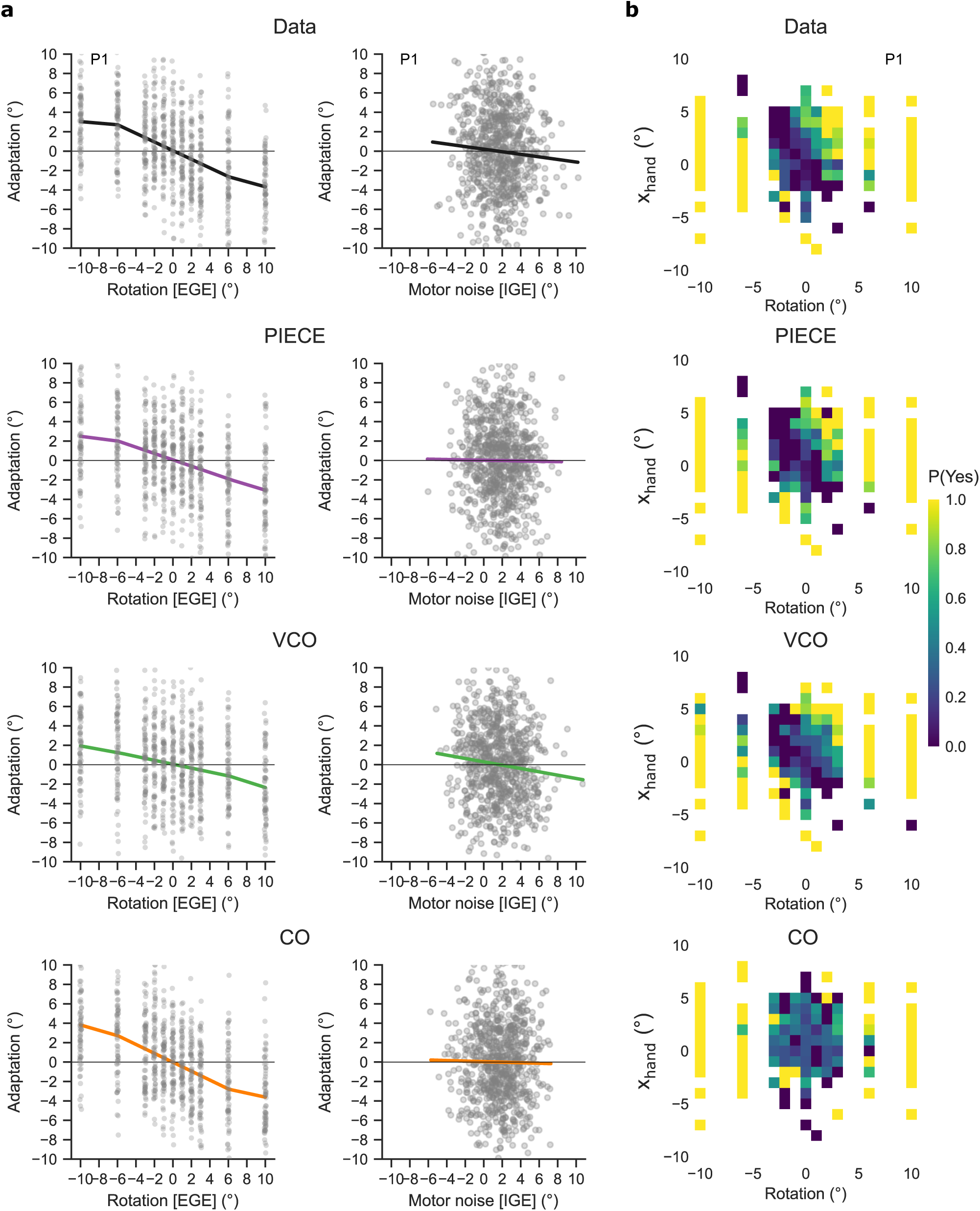
Posterior predictive checks via simulations of both Adaptation and Signal-Detection experiments using maximum-likelihood estimates (MLEs) of each model’s parameter set. Data of a representative participant (Participant 1) on both tasks are shown in the top row. (a) Below the actual data in the top row from P1 on the adaptation task are simulations produced by running each model with its MLEs through the same experimental protocol as P1 experienced. Based on AIC scores, the CO model most accurately captures the differential adaptation to EGE and IGE. As seen in these posterior-predictions, however, there is no stark qualitative difference in predicted behaviors between CO and PIECE (the second most supported model). Notably, the CO model better matches the overall variability of responses demonstrated by P1, whereas PIECE over-predicts response variability relative to what was observed. These patterns were consistently observed across the participants. (b) Below the actual data in the top row from P1 on the signal-detection task are simulations produced by running each model with its MLEs through the same experimental protocol as P1 experienced. Based on AIC scores, the PIECE model best captures the pattern of explicit error detection as a function of perturbation size (x-axis) and actual hand position (y-axis). The posterior-predictions shown here demonstrate PIECE’s alignment with the behavior, predicting a similar band of low-probability “Yes” responses as P1. Dots in (a) represent individual trials.

With regard to participant 14, who appears as somewhat of an outlier based on the AIC scores (Fig 6a), we note that their pattern of adaptation was qualitatively similar to the other participants. Indeed, when removing their data from the adaptation task, the results of our binned analyses (Fig 2) are virtually identical to those when their data are included. The only salient behavioral feature we could detect was that their overall motor output variability increased more during the adaptation block relative to baseline than all participants except for participant 3. As we discuss later, the stochastic nature of error detection during adaptation in VCO and CO models can more readily accommodate such behavioral variability than PIECE can.

**Figure 6:**
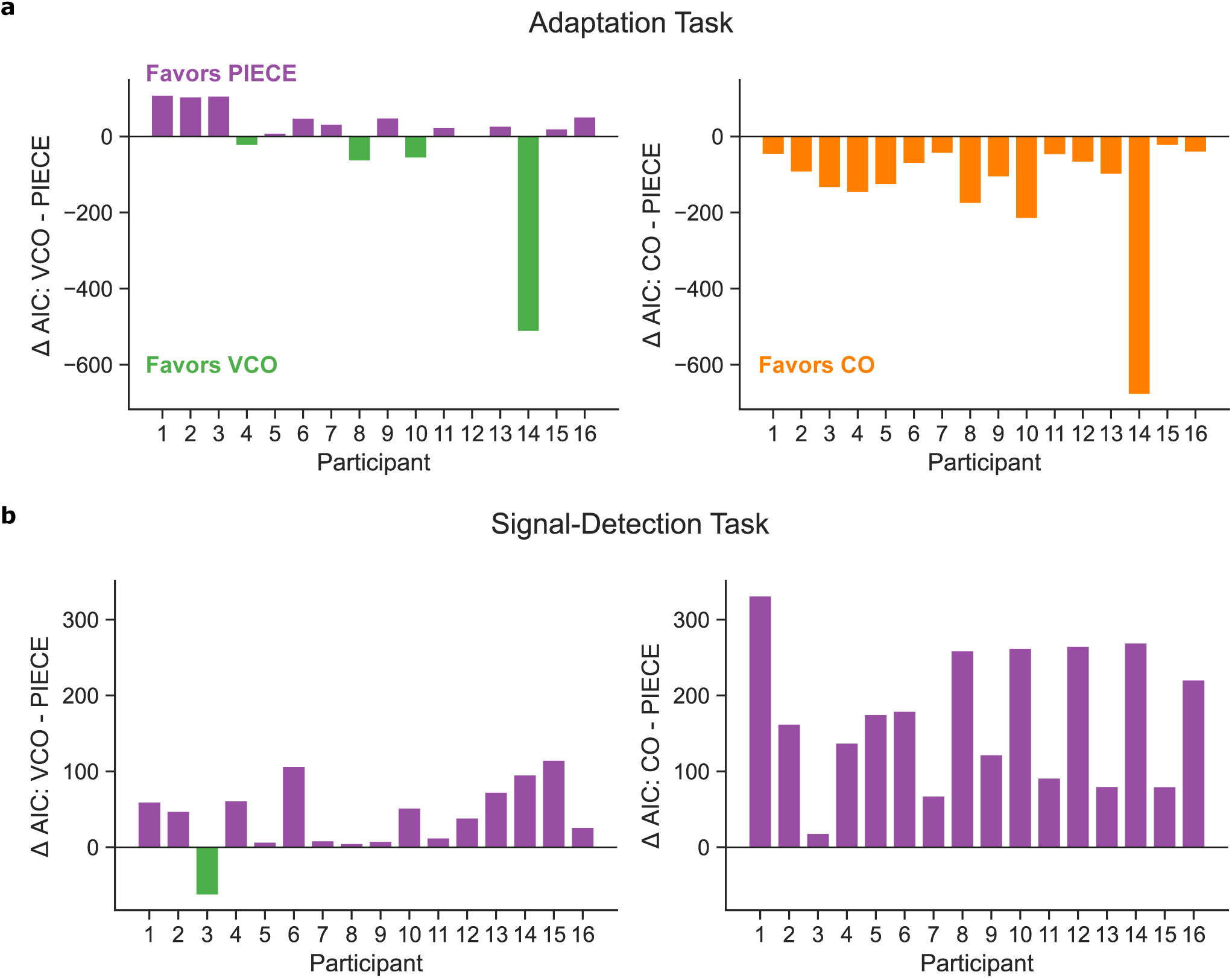
Direct comparisons of AIC scores between VCO and CO models versus PIECE for the (a) adaptation task and the (b) signal-detection task. The bars are color coded to indicate which model is favored for each participant.

For our signal-detection task, the results flipped with PIECE outperforming the VCO and CO models for 15/16 and 16/16 participants, respectively (Fig 6). The CO model performed the worst of the three models, highlighting the dissociation in computational strategies employed by the sensoriomotor system across tasks. The success of PIECE suggests that participants combine prior expectations regarding the perturbation with incoming sensory information from vision and proprioception. This is most clearly visualized by the posterior-predictive heatmaps in Fig 5b, where one can see that the band of lowest probability in the participant’s responses fall somewhere between the vertical column (where rotations are close to zero) predicted by the CO model and the 45° band (where the cursor distance to the target is close to zero) predicted by the VCO model. To summarize the modeling results, of all 9 possible combinations of models across both tasks, for 15 out of 16 participants, the CO and PIECE models best fit adaptation and perturbation behaviors, respectively. One participant’s data was best fit by the combination of CO and VCO.

The use of optimal decision-making in the signal-detection task, in comparison with the more heuristic approach for implicit adaptation, suggests that while the goals in both tasks of estimating, or only detecting, a perturbation requires integrating sensory information, the exact strategies for computing these estimates (or probabilities) are distinct.

## Discussion

Results across our two experiments demonstrate a clear behavioral dissociation between implicit adaptation and explicit perturbation detection. While implicitly adapting to EGEs of 1°, participants could not reliably report the presence of an EGE until it reached ∼ 4°. The CO model unanimously outperformed the PIECE and VCO models on the adaptation task. These findings suggest that when implicitly adapting, the sensorimotor system may rely on a simple “strategy” of subtracting off the estimated reach trajectory (via proprioception and/or efference copy) from the visual cue and correcting for the remaining error. In contrast to this heuristic-like strategy, the superior performance of PIECE on the signal-detection task suggests that participants optimally combine their priors with incoming sensory information when having to make a binary decision regarding the presence of a perturbation. Combined, these results advance our understanding of the computational principles underlying the sensorimotor system’s task-dependent processing of motor errors.

### Implicit movement-error parsing depends on comparing proprioceptive and visual feedback

Our results support the view that the primary computational goal of adaptation is to correct for any external perturbation to our movements, as recent work has emphasized (Kim et al., 2025; Ranjan, 2022). Further, rather than inferring the source of the error and correcting for errors assigned to internal causes, in other words IGE, as has previously been posited (Wei & Körding, 2009), our results show that it is exactly those errors that are thought to have an external, or “world-based”, cause that should be inferred and compensated for, consistent with classic views of motor adaptation (Donchin et al., 2003; Smith et al., 2006; Thoroughman & Shadmehr, 2000). This is the most parsimonious explanation for why we observed robust adaptation to all EGEs, including our smallest perturbation size of 1°, while spontaneously generated IGEs during these same reaches were largely ignored.

Our results are in stark contrast to current computational models of implicit adaptation that posit perceptual errors as the main driver of this learning process (Tsay, Kim, et al., 2022; Zhang et al., 2024). Specifically, according to these models, during a visuomotor rotation paradigm, the percept of hand position is thought to be misaligned with its true position due to sensory conflict between visual and proprioceptive cues. According to these theories, the goal of adaptation is to either realign the “felt” hand position with the target or to minimize this error through state updating. However, as we recently showed, these models do not provide a plausible mechanism for error parsing (Kim et al., 2025). Since the goal in these perceptual error models is to correct for misalignment between felt hand position and the target, obligatory adaptive responses in these models are generated for IGEs as well as EGEs, as missing the target due to motor noise is not treated differently from missing the target due to an EGE. In contrast to perceptual error models, our results suggest that inferences regarding body or limb position should be viewed as subservient to the goal of estimating the state of the perturbation itself.

The observer in the winning CO model compares the discrepancy between visual and proprioceptive cues to an internal threshold, which, when exceeded, leads to a correction that is a fixed proportion of this error. Under the assumptions that proprioception is centered on the actual hand position and vision is centered on the visual cursor position (the sum of hand and experimenter-imposed rotation angles), this reduces to judging whether the noisy estimate of the perturbation itself exceeds the individual participant’s internal threshold (or tolerance) value for visual-proprioceptive mismatches. In sharp distinction to the perceptual-error models, it is the internally measured discrepancy between cues that serves as the error signal, as opposed to the integrated percept of hand position. This model functionally accomplishes the same goals as the Bayesian PIECE model of judging whether a perturbation is present or not and correcting for some proportion of that perturbation. However, an important distinction between PIECE and CO is that the CO model does a simple comparison of cues, whereas the PIECE observer considers all relevant variables within the task to infer both the presence and magnitude of the perturbation. Indeed, the CO model can be viewed as accomplishing the same goals as PIECE in a more direct manner. This apparent preference on the part of the sensorimotor system for a more heuristic strategy when implicitly adapting compared to the computationally more intensive task of making statistically optimal decisions is consistent with other studies of rapid motor decision-making (Adkins et al., 2022), sports performance (Raab, 2017), and sensorimotor behaviors of artificial agents (Batta & Stephens, 2019). Our modeling results align with these studies’ view that when it comes to fast, skillful actions, the sensorimotor system is biased more towards efficiency, in terms of energy or computation, than maximizing expected gain or accuracy. In other words, if the motor system can take a shortcut to doing ‘good enough’, it will (Haith, 2026).

### Bayesian and non-Bayesian accounts of adaptation

Although the CO model convincingly outperformed PIECE based on objective model-selection criteria, we remain somewhat circumspect in our interpretation of these results. The CO model in its current form has difficulty accommodating the saturation of adaptive responses for larger errors that has now been reported in numerous studies, including the present one (Kim et al., 2018; Marko et al., 2012; Morehead et al., 2017). In Fig 5, the CO model shows a slight attenuation of adaptation for the 10° rotation. This is due to the error-detection process involving the comparison of two noisy sensory cues (*v − p*), which can occasionally lead to even larger perturbations occasionally going undetected and therefore resulting in no adaptation on some reaches. In other words, the attenuation at a given perturbation magnitude can result via averaging two categorically different types of responses, adaptive and non-adaptive, due to the stochastic nature of the error and its probabilistic crossing of the detection threshold. (This is, incidentally, also why the CO model is more accommodating to highly variable behavior in model fitting, as we pointed out with respect to participant 14.) As the learning rate in the CO model is a fixed proportion of the error, had we tested a variety of larger perturbation sizes, the lack of attenuation would have been more salient.

PIECE, on the other hand, provides a parsimonious account of the saturation that is based on both the empirical and modeling work of others (Zhang et al., 2024). An earlier study showed that the eyes automatically orient themselves towards the reaching target (Neggers & Bekkering, 2002), while the recent work of Zhang and colleagues (2024) showed that visual uncertainty increases linearly as the visual cursor lands further from the target. For PIECE, this effectively reduces the learning rate, all else being held constant. Similar to a Kalman-filter model (Burge et al., 2008), the effective learning rate in PIECE, which results from the combination of posterior beliefs in the perturbed state and the rotation magnitude, is inversely related to measurement noise. Incorporating increasing visual uncertainty into the CO model could make it appear to accommodate saturated responses for a limited range, but again, only through averaging of a mixture of adaptive and non-adaptive responses. While this provides an interesting alternative explanation for saturated adaptation, we observed no clear evidence of multi-modality in adaptive responses for the larger perturbation sizes, nor has it been reported in other studies utilizing a wider range of perturbations (Tsay, Haith, et al., 2022; Wei & Körding, 2009; Zhang et al., 2024), where the divergence between adaptive and non-adaptive responses would have been most apparent. Furthermore, incorporating increasing visual uncertainty into the CO model would also negatively impact the model’s ability to capture linear adaptive responses to small EGEs, thus hurting its fidelity with respect to adaptation to small perturbations.

Despite these caveats, the CO model clearly did not suffer substantively in the model fits in the present study. This is due to the responses to EGEs up to 6° being highly linear, which the CO model could readily handle. Additionally, only the *±*10° perturbation magnitude, out of the 10 total non-zero rotations, safely fell within the saturated zone (Kim et al., 2018), thus minimizing the leverage of those data points. Indeed, as we pointed out in the Results, fitting a straight line to the binned adaptation versus EGE data only marginally impacted the variance explained as compared to a breakpoint regression (92% vs. 95%). While the PIECE model can more readily accommodate saturated adaptation, it did not do as well in the largely linear zone of adaptive responses. This is because of the complicated interactions between error size, posterior belief in the perturbation (which will generally be lower for smaller perturbations), and estimates of perturbation size, which are, as explained above, impacted by the dynamically changing visual uncertainty. Indeed, when comparing PIECE and CO (and VCO) models on a more constrained adaptation task where rotations fell between *−*4° and +4° as in our prior study (Kim et al., 2025), CO was again the clear winner (S8 Fig).

One additional reason for CO outperforming PIECE is the systematically lower output variability of CO compared to PIECE due to there being fewer sources of uncertainty in the CO model. Most prominently, the CO observer’s learning rate is a fixed proportion of the estimated error size as compared to the PIECE observer, where the effective learning rate is the posterior on Cause, which is a function of noisy sensory measurements that stochastically vary from trial-to-trial. One can see in Fig 5a that the variability of predicted adaptation, relative to the actual data, is slightly exaggerated in the case of PIECE but not for CO. This is borne out in our analysis of the mean-squared errors for model-predicted adaptive responses versus empirically measured adaptation (S7 Fig).

### Future directions for models of implicit adaptation

Overall, the modeling results provide much greater support for application of a simple heuristic than optimal motor decision-making when implicitly adapting. However, one would ideally like to be able to point to qualitative differences in model-predicted outputs. Our parameter- and model-recovery analyses (see S5, S6, and S7 Figs) suggest that there are gains to be made by adjusting the experimental design, as the decision threshold, *D*, was one parameter that was not well-recovered. Although the CO model was the decisive winner when it came to implicit adaptation in our study, for the reasons discussed we do not conclude that it is a comprehensive or “correct” model of adaptation. Future modeling work that combines the heuristic strategy of the CO model with an update rule incorporating other features of the task, rather than the arbitrary constant learning rate currently used in the CO model, will further advance our understanding of the principles underlying the non-linear relationship between EGEs and adaptation.

A unifying model of implicit adaptation must be able to explain a range of sensorimotor learning phenomena. For example, we previously showed that PIECE can readily accommodate the saturation of adaptive responses observed for a wide range of perturbation sizes (Kim et al., 2025). However, future modeling work building off of the present study should address other “standard” implicit adaptation phenomena, such as accumulated adaptation, decay of adaptation during trials with no visual feedback, and responses to clamped visual feedback in which the visual error remains constant (Kim et al., 2018; Morehead et al., 2017).

To model accumulated adaptation in response to a consistent rotation and washout in the absence of visual feedback requires inclusion of a retention factor, the *A* term in a state-space model (or state transition model if working with a Kalman filter) (Haith & Krakauer, 2013; Smith et al., 2006). However, our previous empirical and simulation-based analyses showed that there is little retention in randomized perturbation paradigms and that even a very modest level of retention (e.g., *A* = 0.15) is inconsistent with the error parsing observed across studies (Supplement to Kim et al. 2025). Therefore, to limit the number of parameters and remain consistent with other studies of trial-by-trial adaptation (Burge et al., 2008; Ranjan, 2022; Zhang et al., 2024), we do not include a retention factor in our models. A unifying model of adaptation must capture this dependence of trial-by-trial retention on perturbation statistics.

Another area to explore is how combining ideas across different model architectures may be able to account for not only movement-contingent rotations, as in the current study, but also clamped feedback. Models such as the CO model cannot accommodate adaptation to a visual clamp as the proprioceptive cue remains centered on the actual hand position, which means that the error signal driving adaptation (*v − p*) would increase in an unbounded manner (i.e., *v* stays constant and *p* goes further and further away from *v* as one adapts). However, incorporating cross-modal recalibration such that the observer’s proprioceptive cue is shifted by the visual cue (and vice versa), as we have previously proposed (Tsay, Kim, et al., 2022), is one way to constrain asymptotic adaptation. Interestingly, Bayesian causal inference may provide another interpretation of adaptation to clamped feedback. Despite the experimenter’s usual instructions to “ignore” the visual clamp, the observer may still incorrectly, and implicitly, attribute some degree of belief to the hypothesis that the angular trajectory of the visual error is contingent on their actual reach, driving implicit adaptation. Depending on the dynamics of how those beliefs evolve, it is possible to replicate classic learning curves in response to visuomotor perturbations (Berniker & Körding, 2011), including from clamped feedback.

### Explicit perturbation detection depends on an optimal integration of sensory cues and priors

In contrast to the highly accurate and precise error parsing underlying implicit adaptation, we observed relatively high explicit perturbation-detection thresholds. Consistent with prior work, it required EGEs that were at least 1.5*x* normal baseline motor variability, or ∼ 4°, before participants could report the presence of a perturbation with a performance level of *d^′^* = 1. Further, we showed that on the signal-detection task participants combined prior expectations regarding the perturbation with incoming sensory information in a statistically optimal manner rather than implementing a heuristic, as observed for adaptation. These results point to another clear example where perception for action vs. conscious detection, or recognition, are distinct, as most famously reported for the visual system (Goodale et al., 1986).

In the prior study of explicit perturbation detection by Gaffin-Cahn and colleagues (2019), the authors used a causal-inference model that was structurally identical to PIECE, yet reported that the Visual-Cue-Only (VCO) model was most successful in capturing participants’ detection behavior. In other words, participants in their study based their explicit reports purely on the distance of the feedback cursor from their average reach direction, or motor bias, and whether it crossed an internal detection threshold, with no attention to proprioception. At face value, the clear divergence between the participants’ strategies for signal-detection in the earlier report and the current study is unexpected. However, we note that although PIECE was the best-fit model to 15/16 participants, the VCO model performed second best, and the two models’ predictions were closer to each other than either was to the qualitatively distinct predictions of the CO model. As seen in Fig 5b, the band of low detection probability for VCO is centered on a diagonal that is 45° counter-clockwise of vertical, where the visual error is zero. For the CO model, low probability is centered on the vertical column where the rotation magnitude is zero. Both the behavior for this representative participant and the PIECE predictions are slightly less than 45°, indicating the influence of proprioception on detection judgments. Below we discuss several methodological reasons for why our participants may have been more willing to incorporate a prior and proprioception into their perceptual judgments.

### Strategies for perturbation detection depend on task constraints

We first note that our signal-detection methods aimed to closely match the methodology of our adaptation task, established earlier, in order to confidently make comparisons between the precise error parsing of implicit adaptation to the signal-detection properties of human observers. Thus, online visual feedback was provided during all reaches, as opposed to the endpoint-only feedback used in the Gaffin-Cahn et al. study, to elicit more robust adaptation (Taylor et al., 2014) and maximize the signal-to-noise properties during the adaptation task. We also provided binary feedback only regarding the accuracy of participant judgments, as opposed to veridical feedback regarding the actual hand position at the end of each trial as they did. As Gaffin-Cahn and colleagues noted in their work, providing veridical visual feedback may have biased participants towards prioritizing vision over proprioception. Another difference in methodology was that in our experiment both the reaches and presentation of visual feedback were in parallel horizontal planes, with the visual feedback being provided directly above the stylus position (plus any visual perturbation; parallax difference of ∼ 15cm). In contrast, Gaffin-Cahn and colleagues provided visual feedback on a vertically-oriented monitor while participants made reaches in the horizontal plane. This additional coordinate transformation of sensory signals is an additional source of noise when having to compare proprioceptive and visual cues and may also have caused their participants to be more heavily biased towards using vision (van der Graaff et al., 2017; Wang et al., 2026). Lastly, in our experiment participants made only straight ahead reaches to a single target whereas in the Gaffin-Cahn study visual targets were provided uniformly across the 360° workspace. Previous studies have shown that proprioceptive acuity may increase with repeated reaches (Wong et al., 2011) and is higher in the forward-backward direction than other directions (E. T. Wilson et al., 2010), potentially increasing the likelihood of incorporating proprioception into psychophysical judgments in our experiment. While any one of these methodological differences may not fully explain why participants would ignore proprioception in the Gaffin-Cahn study, they provide multiple plausible reasons proprioception may have been down-weighted as compared to during our task. Regardless of the exact reasons, a parsimonious view suggests that strategies for detecting the presence of a perturbation are more flexible and task-sensitive than the stereotyped operations of the implicit adaptation system (Bond & Taylor, 2015; Morehead et al., 2017).

### The potential functions of high explicit perturbation-detection thresholds

Regardless of how signal-detection was accomplished, both the current study and that of Gaffin-Cahn and colleagues reported high detection thresholds during the task, as compared to the precise error parsing observed in the present study during implicit adaptation. The reason detection thresholds are high relative to the precision of implicit error parsing may lie in their respective functions. Whether we are considering a professional baseball player sending their arm out to catch a ball arriving at 100+ mph or our own rapid text typing on the phone, a sensorimotor miscalibration on the order of a degree can be highly detrimental to performance. There is a growing body of evidence now pointing to this low tolerance for sensorimotor mismatches on the part of the motor system, including several studies of movement error parsing (Kim et al., 2025; Ranjan, 2022; Ranjan & Smith, 2018), adaptation to tiny clamped visual errors, which participants are fully aware of and even told to ignore (Kim et al., 2018, 2019), and adaptation to small amplitude sinusoidal perturbations (Hudson & Landy, 2012). Fine calibration of sensorimotor mappings was likely important for our species’ survival and offloading these functions to systems demanding few cognitive resources is an effective strategy from an evolutionary perspective.

Perhaps more counter-intuitively, less reliable explicit detection of small perturbations may also be beneficial for motor performance. Assuming that perturbation detection precedes generation of an explicit strategy, it could be detrimental to performance to detect tiny errors, as explicit strategies are known to be highly idiosyncratic and noisy, and thus lead to more erratic behavior (McDougle & Taylor, 2019; Taylor et al., 2014). For *de novo* skill learning, this behavioral variability is often viewed as desirable, as it allows an organism to explore more action-based hypotheses (Ding et al., 2026; Tsay et al., 2024). However, for fine tuning of a well-learned movement like reaching, such variable behavior is detrimental to performance (Miyamoto et al., 2020).

### Conclusions

The implicit motor-adaptation system maintains exquisite calibration of sensorimotor mappings by robustly responding to tiny mismatches between visual and proprioceptive cues. Importantly, these adaptive responses correct for the external perturbation and are highly sensitive, as we observed accurate and precise discrimination between internally and externally generated errors as small as 1°. While our results were robust, with consistent behavioral signatures across participants, our sample of 16 neuro-typical young adults, though normal for sensorimotor studies, somewhat limits the generalizability of our findings. Despite this, our modeling analyses further bolster the present findings and suggest that for implicit adaptation, the sensorimotor system employs a simple heuristic of comparing proprioceptive and visual cues to detect and then correct for a fixed fraction of the externally generated error. In contrast to implicit adaptation, reliable explicit perturbation detection did not occur until the perturbation reached ∼ 4°. Also, instead of employing a heuristic strategy, during explicit detection, participants relied on both prior expectations and incoming sensory information in a statistically optimal manner to make their judgments. The combined behavioral and computational modeling results point to a divergence in sensorimotor strategies during perception for action versus perception for recognition. Future work focused on the task-dependent availability and processing of sensory information will contribute to the growing body of work (Haith et al., 2008; Masselink & Lappe, 2021; Simani et al., 2007; Tsay, Kim, et al., 2022) on the organization of perception and skilled action within the nervous system.

## Methods

### Participants

Sixteen healthy, young adults were recruited from the University of British Columbia community (8 females, 8 males, mean age = 21.7; range=[19, 26]). Fifteen of the 16 participants were right-handed according to self-report. All participants were naive to the purpose of the study and provided informed written consent (in accordance with the Declaration of Helsinki) and were remunerated $30 across both testing days. The procedures were approved by the behavioral research ethics board at the University of British Columbia under study ID H23-02324.

### Experimental Set-Up

Participants sat in front of a horizontally-oriented, monitor (144 Hz, 53.2 cm by 30 cm, ASUS), mounted directly above a graphics tablet (49.3 cm by 32.7 cm, Intuos 4XL; Wacom, Vancouver, WA), as shown in Fig 1. Participants performed reaching movements with their dominant arm across the tablet’s surface while holding a stylus embedded within a 3D printed grip. The stylus’s position was recorded at a sampling rate of 200 Hz. The experiment was custom-built via the Psychtoolbox extension in MATLAB (Brainard, 1997).

On each trial, the monitor displayed a yellow start circle (6 mm diameter) positioned in the center of the screen and a green target (6 mm) positioned 9 cm forwards of the start position. The screen also displayed a white cursor (4 mm), representing the stylus’ position. The participants were unable to see their hand or arm, and the experiment was conducted in a dimly-lit room to reduce any additional peripheral vision. In the signal-detection task, participants responded to on-screen text with a mouse controlled with their non-dominant hand.

### General Procedure

Participants completed two sessions on separate days. The order of these sessions was counterbalanced so that half the participants began with the adaptation task and the other half began with the signal-detection task. The purpose of having two separate experimental sessions was to maximize performance on each individual task. That is, we wanted to optimize movement error parsing during the adaptation task and explicit perturbation detection during the signal-detection task.

For both experiments, the participants were instructed to reach as quickly and accurately as they could to the green target, in a single point-to-point movement. To initiate a trial, participants were required to hold their cursor within the start position for 300 ms. Movement time, the time elapsed between movement onset and movement offset, was required to be under 500 ms. Failure to move within this time limit resulted in the green target turning red and reminders from the experimenter to reach more quickly. Movement onset was defined as the first time point at which movement velocity exceeded 1 cm*/*s and the hand had traveled at least 0.5 cm. Movement offset was defined as the first time point after movement onset at which the movement velocity fell below 1 cm/s. The endpoint position was taken at movement offset and endpoint feedback was provided for 500 ms. For the adaptation task, following endpoint feedback, the cursor remained visible throughout the return to the start circle. For the signal-detection task, endpoint feedback was immediately followed by instructions in white text against a black background (see below). Following this, feedback of the cursor was removed until the hand was 1.5 cm away from the start circle, in order to limit the additional time participants had to compare visual feedback of the cursor with proprioceptive information regarding their hand position after the end of the judgment phase. For both tasks, the perturbation, if present, remained on during the search for the start circle, matching our previous methods and prior error-parsing studies (Kim et al., 2025; Ranjan, 2022). Once inside the start circle, the next trial began.

### Adaptation Task

This task quantified single-trial adaptation, utilizing the methods introduced by Ranjan and Smith (2018) and further applied by Kim and colleagues (2025). First, the participants completed 100 baseline trials with no perturbation. This familiarized the participants with the set-up and task, while also enabling quantification of motor noise, or IGE. Immediately prior to the experimental block, in order to discourage the use of explicit strategies, the participants were told that they may notice changes to the cursor feedback, but to continue to aim directly to the target with quick and accurate movements. Perturbation trials (n = 770; 70 trials per rotation amount) had rotations of 0*^◦^*, *±*1*^◦^*, *±*2*^◦^*, *±*3*^◦^*, *±*6*^◦^*, or *±*10*^◦^*. Perturbation trials alternated with null trials (n = 770). Consistent with prior work (Kim et al., 2025; Ranjan & Smith, 2018), half of the null trials (randomly selected) had no visual feedback. Based on our simulations prior to this study (see Supplement in Kim et al., 2025), inclusion of no-visual-feedback trials increases sensitivity to detect adaptation to IGE, if present. In our analyses, the no-visual-feedback trials were treated identically to the null trials with visual feedback. Since the relevant dependent variable is reach angle at maximum radial velocity, which indexes the feedforward motor plan, the presence or absence of visual feedback has no influence. In total, participants executed 1640 reaches across the baseline and experimental blocks. Rest breaks of at least 1 minute duration were provided every 100 trials.

### Signal-Detection Task

This task assessed participants’ ability to detect visual perturbations. As in the adaptation task, participants executed 100 baseline reaches. To acquaint participants with visual perturbations and the task, they completed six instruction trials. Instruction trials consisted of two null trials and perturbations of *±*2° and *±*10°, providing participants with examples of EGE magnitudes that were below and above previously reported detection thresholds (Gaffin-Cahn et al., 2019). Following each reach, participants were asked to report whether they detected the presence of a perturbation. They did so by clicking either the left mouse button for ‘yes’ or the right mouse button for ‘no’. Following their initial answer, participants had the option to confirm or change their response. This option was provided to allow participants to correct for errant key presses. Following their final answer, binary feedback appeared on screen, indicating the correctness of the response (Fig 1). During the instruction trials, and following binary feedback, visual feedback of the target, the rotated cursor, and the actual hand positions were also provided.

Prior to the start of the experimental block, participants were informed that half of the trials had a perturbation. Participants experienced the exact same perturbation sizes as in the adaptation task. Unlike the adaptation task, however, every trial had visual feedback and the scheduling of perturbations was randomized in order to remove any predictability to the trial schedule. The psychophysical judgment phase of each trial was identical in structure to the instruction block except that participants no longer received additional veridical visual feedback of their hand position after making their decision. These experimental design decisions—explicitly informing participants of the proportion of null trials, randomizing the perturbation schedule, and providing binary feedback after each judgment—helped to optimize participants’ perturbation-detection performance (Landy, 2023). However, it also meant that participants had explicit knowledge of whether they were perturbed or not following each reach. Because of this and the following two reasons, we made the *a priori* decision to not focus our analysis on implicit adaptation during this task. The randomization of the perturbations negated our ability to measure movement error parsing (Kim et al., 2025; Ranjan, 2022). Also, the psychophysical judgment and binary feedback phases substantially increases the intertrial interval as compared to the adaptation task (by a factor of 2.3), thus attenuating implicit adaptation (Hadjiosif et al., 2023).

In total, the experimental block was composed of 800 trials (400 null; 400 perturbation; 40 per non-zero perturbation level). In total, there were 906 trials across the baseline, instruction, and experimental blocks. Rest breaks of at least 1 minute duration were provided every 100 trials.

### Data Analysis

All analyses shown in this document were conducted using custom-written Python scripts using standard Python libraries: NumPy (Harris et al., 2020), Pandas (Team, 2020), SciPy (Virtanen et al., 2020), Matplotlib (Hunter, 2007), and Seaborn (Waskom, 2021). Reach angle was defined as the angle between straight lines connecting the start position to the target and the hand at peak velocity. For each participant, trials with reach angles exceeding 22.5° were excluded, and then based on the remaining trials, trials with reach angles exceeding a *z*-score magnitude of 3.5 were excluded. For the adaptation task, the average percentage of outlier trials across participants was 0.78% (min: 0.12%; max: 6.79%). Inclusion/exclusion of the only participant who had greater than 1% of outlier trials (participant 3) did not qualitatively change any of the reported results. For the signal-detection task, the average percentage of outliers was 0.49%, (min: 0%; max: 2.5%). Over 99.3% of all experimental trials were retained.

Adaptation was quantified as the difference in reach angles, *x*_hand_, surrounding a perturbation trial: adaptation*_n_* = *x*_hand, n+1_ *− x*_hand, n-1_, where *n* indexes a perturbation trial. Quantifying adaptation in this manner rather than reach*_n_*_+1_ *−* reach*_n_*, as done in many previous studies of single-trial adaptation (Tsay, Haith, et al., 2022; Wei & Körding, 2009), provides an uncontaminated measure of adaptation to internally-generated motor noise (IGE), as it avoids the outcome variable (adaptation) being defined in part by the predictor variable IGE (*x*_hand_), thus creating spurious correlations.

For our signal-detection analyses, we first categorized participants’ responses as either hits (correctly identifying a perturbation) or false alarms (incorrectly identifying a perturbation when none occurred). We applied a standard correction for hit and false alarm rates of 0 and 1 (Hautus, 1995). These rates were then converted into standardized *z*-scores. The detection threshold, formally referred to as the discriminability index (*d′*), is calculated as the difference between the *z*-scores of the hit and false-alarm rates. We define *d′* of 1 as the threshold for reliable perturbation detection, corresponding to 69% correct responses (for a neutral decision criterion).

### Statistical Analyses

All regression and *t*-tests were conducted after visually confirming normality of the data via QQ plots. We initially fit the explicit perturbation-detection data with a logarithmic function (based on (Gaffin-Cahn et al., 2019)), but after seeing the data much more closely resembled a straight line, we switched to linear-regression fits. The improvement in fits was confirmed through comparison of AIC/BIC scores.

## Modeling

Three models were fit to all individual participant data. Two of the models, the Visual-Comparison-Observer and the Comparison-Observer models, were adapted from a previous paper (Gaffin-Cahn et al., 2019). These models were originally developed to account for explicit detection and so were modified here in order to encompass adaptation as well. The third model is the Parsing of Internal and External Causes of Error (PIECE) model, a Bayesian causal-inference model. As described in the Results, for each individual participant, each of the three models was fit simultaneously to data from both tasks as well as separately to the data from the two tasks.

Maximum-likelihood estimation was performed using Bayesian adaptive direct search (*BADS* ; Acerbi and Ma, 2017), via the PyBADS software (Singh & Acerbi, 2024). PyBADS alternates between a series of fast, local optimization steps and a systematic, slower exploration of a mesh grid. The goal of our fitting procedure was to find the parameter set, denoted Θ*_model_*, that minimized the negative log-likelihood of the recorded trial-by-trial data. Consistent with prior studies of trial-by-trial adaptation (Wei & Körding, 2009; Zhang et al., 2024), and based on our finding of minimal residual adaptation between trials during randomized perturbation paradigms (see Supplement in Kim et al., 2025), we assumed trials were independent of each other and therefore summed the negative log-likelihoods across all trials. We used closed-form solutions to compute the log-likelihood in the case of the VCO and CO models for the signal-detection task. For all other cases, we used a minimum of 700,000 Monte Carlo simulations per participant per task to generate noisy sensory cues (as their values are unknown to the experimenter) to closely approximate response distributions on each trial and compute the log-likelihoods. When fitting simultaneously to both tasks, our fitting procedure aimed to minimize the mean of the average per-trial log-likelihood for each task in order to ensure that the tasks were equally weighted. Without this adjustment, the adaptation task would have exercised much greater influence on the parameter estimates due to it having nearly two times as many trials (and thus contributing a much greater proportion than the signal-detection task to the sum of log-likelihoods).

For each participant’s data set, we used five random initial starting points to avoid local minima. Model comparison was completed using Akaike Information Criterion (AIC) scores (Akaike, 2025).

### Parameter and Model Recovery

We performed model- and parameter-recovery analyses with all three models on both tasks to ensure the validity of model comparisons (R. C. Wilson & Collins, 2019) and parameter estimates (S4 Fig), particularly of our winning models. The results of these analyses can be found in the Supporting Information (S4, S5, and S6 Figs) We note here that model recovery for all three models and for both tasks approached 100% identifiability.

### PIECE

The PIECE model (Fig 4) frames error detection and motor adaptation as Bayesian causal inference. The causal node, *C*, gives rise to two competing hypotheses for the observer to discriminate. The first is that the sensory cues are more consistent with an unperturbed world state, meaning any detected movement errors are the result of internal processes. The competing hypothesis is that the sensory cues are more consistent with a perturbed world state.

PIECE posits that the observer optimally integrates two sources of sensory information: vision (*x_v_*) and proprioception (*x_p_*). The visual cue follows a normal distribution and is centered on the cursor position such that 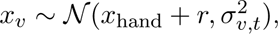 where *r* represents the unknown (to the observer) visuomotor rotation.

Along with proprioceptive information from joint receptors and muscles, *x_p_* is assumed to also incorporate an efference-copy-based motor prediction of the hand’s true position. However, from the experimenter’s perspective, there is no way to dissociate the sensor-based proprioceptive information and motor prediction since they are both centered on the actual hand position. Therefore, *x_p_* is interpreted as encompassing both contributions to felt hand position. The (combined) proprioceptive measurement is normally distributed: 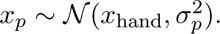

The prior on hand position, *x*_hand_, is normally distributed: 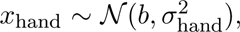 where *b* captures any bias in reach angle and *σ*_hand_ represents motor variability. The observer also has a prior on the rotation magnitude, 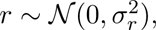 where *σ_r_* quantifies a plausible range of perturbation sizes.

To determine the cause of movement-related error, the PIECE observer applies Bayes’ Rule to form a posterior on *C* as follows:

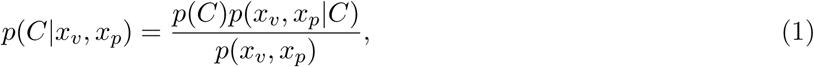

where *p*(*C*) is the prior belief in Cause, and *p*(*x_v_, x_p_|C*) is the joint likelihood function over all sensory cues.

Given that the probability of a perturbation was 0.5, and to constrain the model by avoiding incorporation of an additional free parameter, we assume the observer assigns equal probability to perturbed and unperturbed world states (i.e., *p*(*C*) = 0.5).

Calculating the posterior probability of the unperturbed world state requires marginalizing over *x*_hand_:

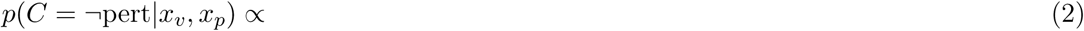

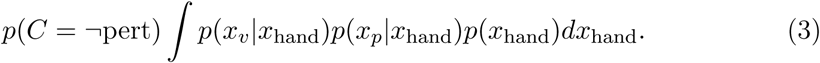

To compute the probability of the perturbed world state the observer must integrate over both *x*_hand_ and *r*:

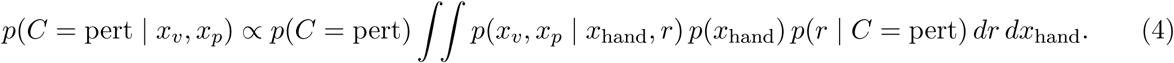

This can be simplified to the final expression shown below:

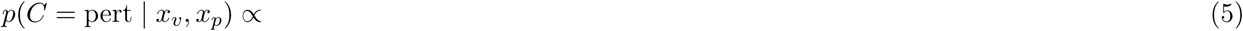

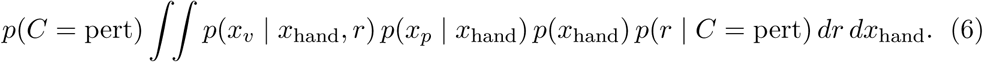

#### Adaptation

Since the true rotation magnitude is unknown to the PIECE observer, they must infer it in order to guide their next reach. Computing a posterior over *r* requires a marginalization over *x*_hand_, as the visual cue is dependent on both the rotation and the actual hand position:

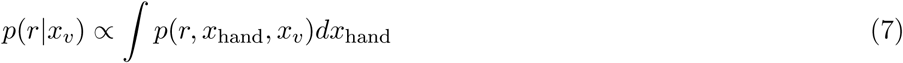

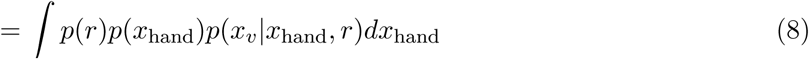

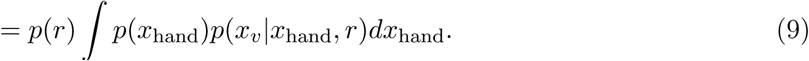

The observer’s estimate of the rotation, *r*^ is the maximum a posteriori (MAP) estimate of *r*, that is, the value of *r* that maximizes *p*(*r|x_v_*). For adaptation, the posterior on cause, *C*, acts as a weight on *r*^, effectively serving as a learning rate that is derived from the statistics of the task, as opposed to being a free parameter as in the other two models. We assume the observer’s motor output is a reflection of this estimate (thus opposite-signed), with the addition of the observer’s motor bias and intrinsic motor noise.

Thus, hand position on trial *t* + 1 is defined as:

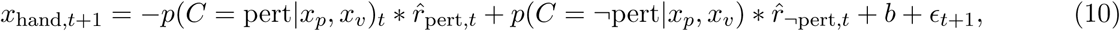

where 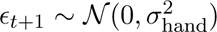 represents motor noise on trial *t*. By definition, in the unperturbed world state, *r*^*_¬_*_pert_*_,t_* is always zero, so the above equation simplifies to:

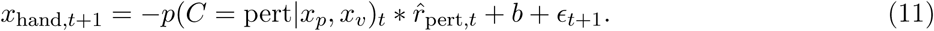

For the adaptation and signal-detection data, PIECE was fit with 5 free parameters: *σ_p_*, *σ_pert_*, *b* and two parameters, *α* and *β*, for fitting the linearly increasing uncertainty in vision as a function of cursor distance to target (Kim et al., 2025; Zhang et al., 2024):

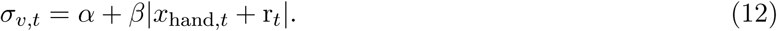

#### Signal Detection

PIECE detects an error once the posterior on cause (*C*) for the perturbed world state is greater than the unperturbed world state. Specifically, if *p*(*C* = perturbed*|x_v_, x_p_*) *>* 0.5, the model predicts a ‘Yes’ response. Otherwise, the model predicts a ‘No’ response.

For PIECE and the other two models, fitting the data with and without a fixed 2% lapse rate (Kingdom & Prins, 2016) had no impact on the results, with both methods generating the exact same proportions of participants favored by each model and nearly identical AIC scores. The lack of impact on our results of including a lapse rate may be because participants were allowed to correct for errant key presses and their explicit reports regarding the visual perturbation were made immediately following visually guided reaches to a clear visual target, making attention to the sensory stimuli compulsory. Therefore, we report the more parsimonious models fits that do not include a lapse rate.

### Visual-Cue-Only Observer

#### Signal Detection

The Visual-Cue-Only (VCO) Observer relies solely on visual feedback to detect a perturbation. This model assumes the observer compares a noisy perceptual estimate of the cursor’s final position, *x_v_*, to the expected endpoint, represented by *b*, the observer’s intrinsic motor bias. These sensory estimates are drawn from normal distributions:

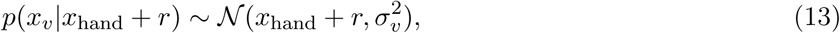

where *x*_hand_ is the final hand position, *r* is an applied visual perturbation, and 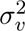 is visual uncertainty. The observer computes the difference between the perceived cursor location and the observer’s intrinsic bias, *e_t_* = *x_v_ − b*. A perturbation is detected when the magnitude of the difference exceeds a decision threshold, that is, when

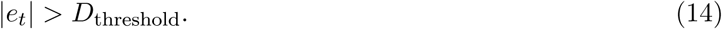

*D*_threshold_ thus represents the discrepancy between the visual cue and motor bias for which the probability of detection is 50%.

We note that our original intention in this study was to implement the VCO and CO models in a manner consistent with Gaffin-Cahn and colleagues (2019). However, we also implemented variants of the VCO model and CO models that incorporated increasing visual uncertainty (Eqn. 12), as we do for PIECE. In both cases, this modification either worsened the quality of fits for a majority of participants, as judged by AIC scores (VCO), or had no qualitative effect on the results (CO). We have therefore chosen to focus our reporting on the more constrained implementation of the models as described above.

#### Adaptation

To accommodate adaptation, we assumed that the VCO observer corrects for a proportion of the detected error the following reach, 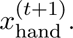 If no error is detected, no learning occurs:

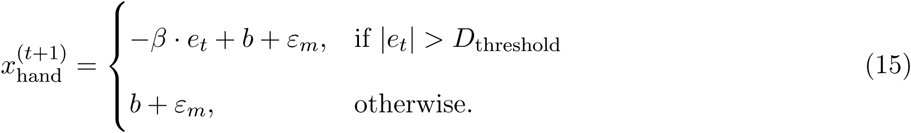

There is a total of four free parameters in the VCO model : *D*_threshold_, *b*, *σ_v_*, and *β*. However, *β* was not used for the signal-detection task since we were only concerned with fitting the psychophysical judgments in that experiment.

### Comparison Observer

#### Signal Detection

The Comparison Observer (CO) makes a decision regarding the presence of a perturbation by comparing visual and proprioceptive cues. A perturbation is detected if the separation between sensory cues is sufficiently large, that is, when

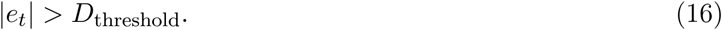

Since the mean of *x_v_* is the cursor position and the mean of *x_p_* is the actual hand position, the discrepancy between the two cues has mean magnitude of *r* and variance 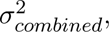 which represents the sum of the visual and proprioceptive variances. We fit 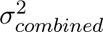 here because the individual contributions of proprioceptive and visual noise are not discriminable in the data.

#### Adaptation

As with the VCO model, the CO model makes a correction in the subsequent reach that is proportional to the detected error (*e_t_* above):

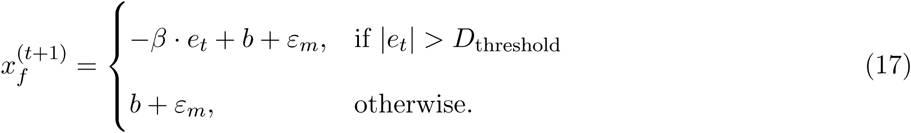

Both *β* and *ε_m_* are, again, the learning rate and motor noise, respectively, with 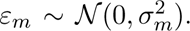 There is a total of four free parameters in the CO model: *D*_threshold_, *b*, *σ_combined_*, and *β*. However, *β* and the motor bias, *b* were not used for the signal-detection task. In the case of the former, we were only fitting the psychophysical judgments, and for the latter, bias plays no role in judging the discrepancy between *x_v_* and *x_p_*.

## Acknowledgments

HEK is funded by a Natural Sciences and Engineering Research Council of Canada (NSERC) Discovery Grant (RGPIN-2025-04104) and a Canada Foundation for Innovation John R. Evans Leaders Fund grant. MSL is funded by NIH grant EY08266. RC is funded by a NSERC Discovery Grant (RGPIN-2026-04716).

## Supporting Information

**S1 Fig. Plots of individual data of all 16 participants from the adaptation task.**

**S2 Fig. Plots of individual data of all 16 participants from the signal-detection task.**

**S3 Table. Maximum-likelihood estimates of each model.** Top table shows maximum-likelihood estimates of each model parameter during the adaptation task. Note: *β* in PIECE refers to the slope associated with increasing visual uncertainty whereas *β* in the VCO and CO models refer to the learning rate for adaptation. Bottom table shows maximum-likelihood estimates of each model parameter during the signal-detection task.

**S4 Fig. Parameter recovery for the winning CO model in the adaptation task.** To validate the parameter values obtained by fitting CO or PIECE (i.e., the winning models for each task) to our behavioral data and our ability to distinguish between each of the three models, we performed parameter- and model-recovery analyses (R. C. Wilson and Collins, 2019). We generated 100 synthetic datasets with each of the three models used in this study, totaling 300 datasets in all. The synthetic data from each model were generated using parameter values drawn from uniform distributions bounded by the minimum and maximum parameter values obtained from fits to each individual participant’s behavioral data (i.e., the range of maximum-likelihood estimates of each model parameter).

Scatterplots show the CO model parameter values that were recovered (*y*-axes) from fits to 100 synthetic datasets generated with the CO model. The parameter values used to generate the synthetic data are shown on the *x*-axes. Except for the detection threshold, *D*, the recovered parameters for the CO model were all highly correlated with the parameter values used to generate the synthetic data (Pearson’s *r >*= 0.953, *p <* 10*^−^*^6^). In the case of *D*, the correlation coefficient, *r*, was weaker (*r* = 0.248, *p* = 0.013). Future experiments that test a wider range of perturbation sizes will help achieve more accurate recovery of this parameter, as the current experiment spanned a relatively narrow range.

**S5 Fig. Parameter recovery for the winning PIECE model in the signal-detection task.** Scatter plots show the PIECE model parameter values that were recovered (*y*-axes) from fits to 100 synthetic datasets generated with the PIECE model. The parameter values used to generate the synthetic data are shown on the *x*-axes. The recovered parameters for the PIECE model were also, in general, tightly correlated with the parameter values used for simulation. The correlation coefficients, *r*, were all greater than 0.835, except in the case of *β*, the parameter representing the rate at which visual uncertainty increases as a function of visual cue distance from the target, which had a more moderate value of 0.493.

**S6 Fig. Confusion matrices for both tasks.** The value in each cell of the matrix indicates *p*(recovered model | simulated model). We fit all 300 synthetic datasets with each model using maximum-likelihood estimation and performed objective model selection for each dataset. Our model-recovery analysis showed that we can accurately identify which of the three models was responsible for generating each synthetic data set with at least 95% accuracy. And in the case of the PIECE model, 100% accuracy across both tasks. This points to the robustness of our modeling results, as well as the strength of experimental design as we were successfully able to identify the model that generated the data in nearly all cases.

**S7 Fig. Bias-variance decomposition of mean-squared errors (MSE) of model predictions relative to actual data.** Scatterplots of model-predicted adaptation vs. actual adaptation. We use the posterior predictions from Fig 5 and plot their relationship to the actual adaptation produced by P1. The black dashed line represents the unity line, which all dots would fall on if adaptation was perfectly predicted by the model. The red dashed lines represent the line of best-fit through data. The CO model had the lowest mean-squared error (MSE), which means its predicted responses were most closely aligned with actual behavior. This result can be understood as resulting from both a lower bias (i.e., the slope of the best-fit line is closest to 1 compared to the other two models) and lower output variability as compared to the other two models.

**S8 Fig. Model-based analyses of data from Kim et al. (2025).** Results of fitting the three computational models from the current study to data from a prior study of movement error parsing. In our prior study we tested movement error parsing in response to visuomotor rotations of 0°, *±*2°, and *±*4° only. The model selection results here largely mimic those of the present study. All 16 participants’ data were best fit by the CO model, with PIECE performing second best, and VCO performing the worst.

## References

Acerbi, L., & Ma, W. J. (2017). Practical Bayesian optimization for model fitting with Bayesian adaptive direct search. Advances in Neural Information Processing Systems, 30, 1834–1844.

Adkins, T., Lewis, R., & Lee, T. (2022). Heuristics contribute to sensorimotor decision-making under risk. Psychonomic Bulletin & Review, 29 (1), 145–158.

Akaike, H. (2025). Akaike’s information criterion. In M. Lovric (Ed.), International encyclopedia of statistical science. Springer-Verlag.

Batta, E., & Stephens, C. (2019). Heuristics as decision-making habits of autonomous sensorimotor agents. Artificial Life Conference Proceedings.

Berniker, M., & Körding, K. P. (2011). Estimating the relevance of world disturbances to explain savings, interference and long-term motor adaptation effects. PLoS Computational Biology, 7 (10), e1002210.

Bond, K. M., & Taylor, J. A. (2015). Flexible explicit but rigid implicit learning in a visuomotor adaptation task. Journal of Neurophysiology, 113 (10), 3836–3849.

Brainard, D. H. (1997). The psychophysics toolbox. Spatial Vision, 10 (4), 433–436.

Burge, J., Ernst, M. O., & Banks, M. S. (2008). The statistical determinants of adaptation rate in human reaching. Journal of Vision, 8 (4), 20–20.

Carriot, J., Brooks, J. X., & Cullen, K. E. (2013). Multimodal integration of self-motion cues in the vestibular system: Active versus passive translations. Journal of Neuroscience, 33 (50), 19555–19566.

Ding, W., Niyogi, A., Taylor, J. A., & Tsay, J. S. (2026). Hypothesis testing governs strategic motor learning. npj Science of Learning.

Donchin, O., Francis, J. T., & Shadmehr, R. (2003). Quantifying generalization from trial-by-trial behavior of adaptive systems that learn with basis functions: Theory and experiments in human motor control. Journal of Neuroscience, 23 (27), 9032–9045.

Faisal, A. A., Selen, L. P., & Wolpert, D. M. (2008). Noise in the nervous system. Nature Reviews Neuroscience, 9 (4), 292–303.

Gaffin-Cahn, E., Hudson, T. E., & Landy, M. S. (2019). Did I do that? detecting a perturbation to visual feedback in a reaching task. Journal of Vision, 19 (1), 5.

Goodale, M. A., Pelisson, D., & Prablanc, C. (1986). Large adjustments in visually guided reaching do not depend on vision of the hand or perception of target displacement. Nature, 320 (6064), 748–750.

Hadjiosif, A. M., Morehead, J. R., & Smith, M. A. (2023). A double dissociation between savings and long-term memory in motor learning. PLoS Biology, 21 (4), e3001799.

Haith, A. M. (2026). Policy-gradient reinforcement learning as a general theory of practice-based motor skill learning. bioRxiv.

Haith, A. M., Jackson, C. P., Miall, R. C., & Vijayakumar, S. (2008). Unifying the sensory and motor components of sensorimotor adaptation. Neural Information Processing Systems Proceedings.

Haith, A. M., & Krakauer, J. W. (2013). Theoretical models of motor control and motor learning. In Routledge handbook of motor control and motor learning (pp. 16–37). Routledge.

Harris, C. R., Millman, K. J., van der Walt, S. J., Gommers, R., Virtanen, P., Cournapeau, D., Wieser, E., Taylor, J., Berg, S., Smith, N. J., Kern, R., Picus, M., Hoyer, S., van Kerkwijk, M. H., Brett, M., Haldane, A., del Río, J. F., Wiebe, M., Peterson, P., . . . Oliphant, T. E. (2020). Array programming with NumPy. Nature, 585 (7825), 357–362.

Hautus, M. J. (1995). Corrections for extreme proportions and their biasing effects on estimated values of d’. Behavior Research Methods, Instruments, & Computers, 27 (1), 46–51.

Hudson, T. E., & Landy, M. S. (2012). Measuring adaptation with a sinusoidal perturbation function. Journal of Neuroscience Methods, 208 (1), 48–58.

Hunter, J. D. (2007). Matplotlib: A 2d graphics environment. Computing in Science & Engineering, 9 (3), 90–95.

Hutter, S. A., & Taylor, J. A. (2018). Relative sensitivity of explicit reaiming and implicit motor adaptation. Journal of Neurophysiology, 120 (5), 2640–2648.

Kim, H. E., Avraham, G., & Ivry, R. B. (2021). The psychology of reaching: Action selection, movement implementation, and sensorimotor learning. Annual Review of Psychology, 72 (1), 61–95.

Kim, H. E., Chua, R., & Hu, D. (2025). Causal inference, prediction and state estimation in sensorimotor learning. Proceedings of the Royal Society B: Biological Sciences, 292 (2052), 20251320.

Kim, H. E., Morehead, J. R., Parvin, D. E., Moazzezi, R., & Ivry, R. B. (2018). Invariant errors reveal limitations in motor correction rather than constraints on error sensitivity. Communications Biology, 1 (1), 19.

Kim, H. E., Parvin, D. E., & Ivry, R. B. (2019). The influence of task outcome on implicit motor learning. eLife, 8, e39882.

Kingdom, F. A., & Prins, N. (2016). Psychophysics: A practical introduction. Elsevier.

Landy, M. S. (2023). Signal detection theory. Cambridge handbook of research methods and statistics for the social and behavioral sciences, 3.

Marko, M. K., Haith, A. M., Harran, M. D., & Shadmehr, R. (2012). Sensitivity to prediction error in reach adaptation. Journal of Neurophysiology, 108 (6), 1752–1763.

Masselink, J., & Lappe, M. (2021). Visuomotor learning from postdictive motor error. Elife, 10, e64278.

McDougle, S. D., & Taylor, J. A. (2019). Dissociable cognitive strategies for sensorimotor learning. Nature Communications, 10 (1).

Miyamoto, Y., Wang, S., & Smith, M. (2020). Implicit adaptation compensates for erratic explicit strategy in human motor learning. Nature Neuroscience, 23 (3), 443–455.

Morehead, J. R., Taylor, J. A., Parvin, D. E., & Ivry, R. B. (2017). Characteristics of implicit sensorimotor adaptation revealed by task-irrelevant clamped feedback. Journal of Cognitive Neuroscience, 29 (6), 1061–1074.

Neggers, S. F., & Bekkering, H. (2002). Coordinated control of eye and hand movements in dynamic reaching. Human Movement Science, 21 (3), 37–64.

Raab, M. (2017). Motor heuristics and embodied choices: How to choose and act. Current Opinion in Psychology, 16, 34–37.

Ranjan, T. (2022). Understanding the role of internal predictions in sensorimotor adaptation [Doctoral dissertation, Harvard University].

Ranjan, T., & Smith, M. A. (2018). Cancellation of internally-generated errors from the signal driving motor adaptation. Advances in Motor Learning and Motor Control (MLMC).

Simani, M. C., McGuire, L. M. M., & Sabes, P. N. (2007). Visual-shift adaptation is composed of separable sensory and task-dependent effects. Journal of Neurophysiology, 98 (5), 2827–2841.

Singh, G. S., & Acerbi, L. (2024). PyBADS: Fast and robust black-box optimization in Python. Journal of Open Source Software, 9 (94), 5694.

Smith, M. A., Ghazizadeh, A., & Shadmehr, R. (2006). Interacting adaptive processes with different timescales underlie short-term motor learning. PLoS Biology, 4 (6), e179.

Sommer, M. A., & Wurtz, R. H. (2008). Brain circuits for the internal monitoring of movements. Annu. Rev. Neurosci., 31 (1), 317–338.

Szarka, A., Kim, H. E., Inglis, J. T., & Chua, R. (2025). Evidence for an efferent-based prediction contributing to implicit motor adaptation. PLoS One.

Taylor, J. A., Krakauer, J. W., & Ivry, R. B. (2014). Explicit and implicit contributions to learning in a sensorimotor adaptation task. Journal of Neuroscience, 34 (8), 3023–3032.

Team, T. P. D. (2020, February). *Pandas-dev/pandas: Pandas* (Version latest). Zenodo.

Thoroughman, K. A., & Shadmehr, R. (2000). Learning of action through adaptive combination of motor primitives. Nature, 407 (6805), 742–747.

Tsay, J. S., Haith, A. M., Ivry, R. B., & Kim, H. E. (2022). Interactions between sensory prediction error and task error during implicit motor learning. PLoS Computational Biology, 18 (3), e1010005.

Tsay, J. S., Kim, H. E., Haith, A. M., & Ivry, R. B. (2022). Understanding implicit sensorimotor adaptation as a process of proprioceptive re-alignment. eLife, 11.

Tsay, J. S., Kim, H. E., McDougle, S. D., Taylor, J. A., Haith, A. M., Avraham, G., Krakauer, J. W., Collins, A. G. E., & Ivry, R. B. (2024). Fundamental processes in sensorimotor learning: Reasoning, refinement, and retrieval. eLife, 13, e91839.

van Beers, R. J. (2009). Motor learning is optimally tuned to the properties of motor noise. Neuron, 63 (3), 406–417.

van der Graaff, M. C. W., Brenner, E., & Smeets, J. B. J. (2017). Vector and position coding in goal-directed movements. Experimental Brain Research, 235 (3), 681–689.

Virtanen, P., Gommers, R., Oliphant, T. E., Haberland, M., Reddy, T., Cournapeau, D., Burovski, E., Peterson, P., Weckesser, W., Bright, J., et al. (2020). Scipy 1.0: Fundamental algorithms for scientific computing in python. Nature Methods, 17 (3), 261–272.

Wang, T., Morehead, J. R., Jiang, A., Ivry, R. B., & Tsay, J. S. (2026). Motor biases reflect a misalignment between visual and proprioceptive reference frames. eLife, 13, RP100715.

Waskom, M. L. (2021). Seaborn: Statistical data visualization. Journal of Open Source Software, 6 (60), 3021.

Wei, K., & Körding, K. (2009). Relevance of error: What drives motor adaptation? Journal of Neurophysiology, 101 (2), 655–664.

Wilson, E. T., Wong, J., & Gribble, P. L. (2010). Mapping proprioception across a 2d horizontal workspace. PloS One, 5 (7), e11851.

Wilson, R. C., & Collins, A. G. (2019). Ten simple rules for the computational modeling of behavioral data. eLife, 8, e49547.

Wong, J. D., Wilson, E. T., & Gribble, P. L. (2011). Spatially selective enhancement of proprioceptive acuity following motor learning. Journal of Neurophysiology, 105 (5), 2512–2521.

Zhang, Z., Wang, H., Zhang, T., Nie, Z., & Wei, K. (2024). Perceptual error based on bayesian cue combination drives implicit motor adaptation. eLife, 13.

