## Supplementary figures and images for "Distinct computations underlie movement error parsing during implicit motor adaptation and perturbation detection"

### S1 Figure

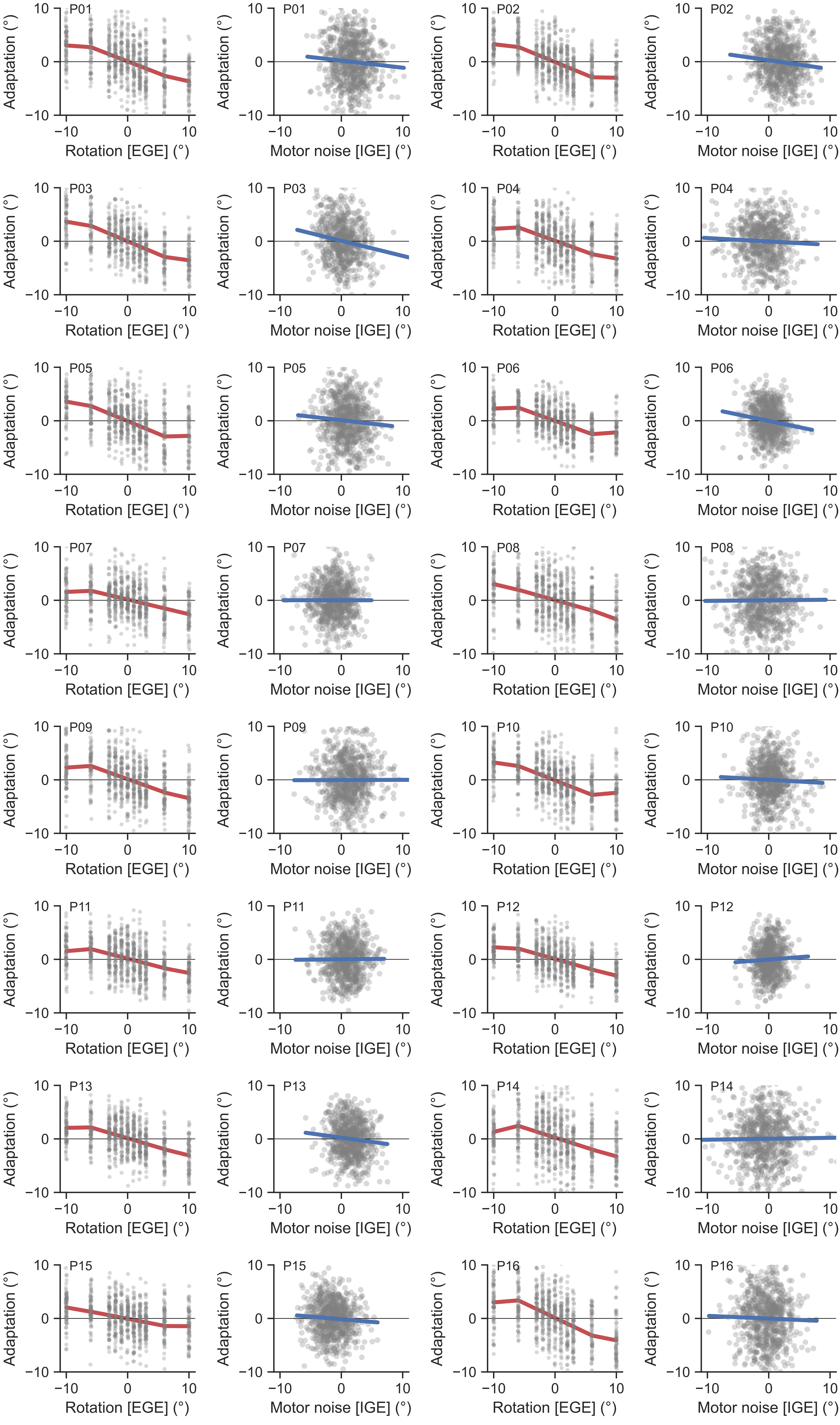
P01
P02
P03
P04
P05
P06
P07
P08
P09
P10
P11
P12
P13
P14
P15
P16
Adaptation (°)
Rotation [EGE] (°)
Motor noise [IGE] (°)
10
0
-10

### S2 Figure

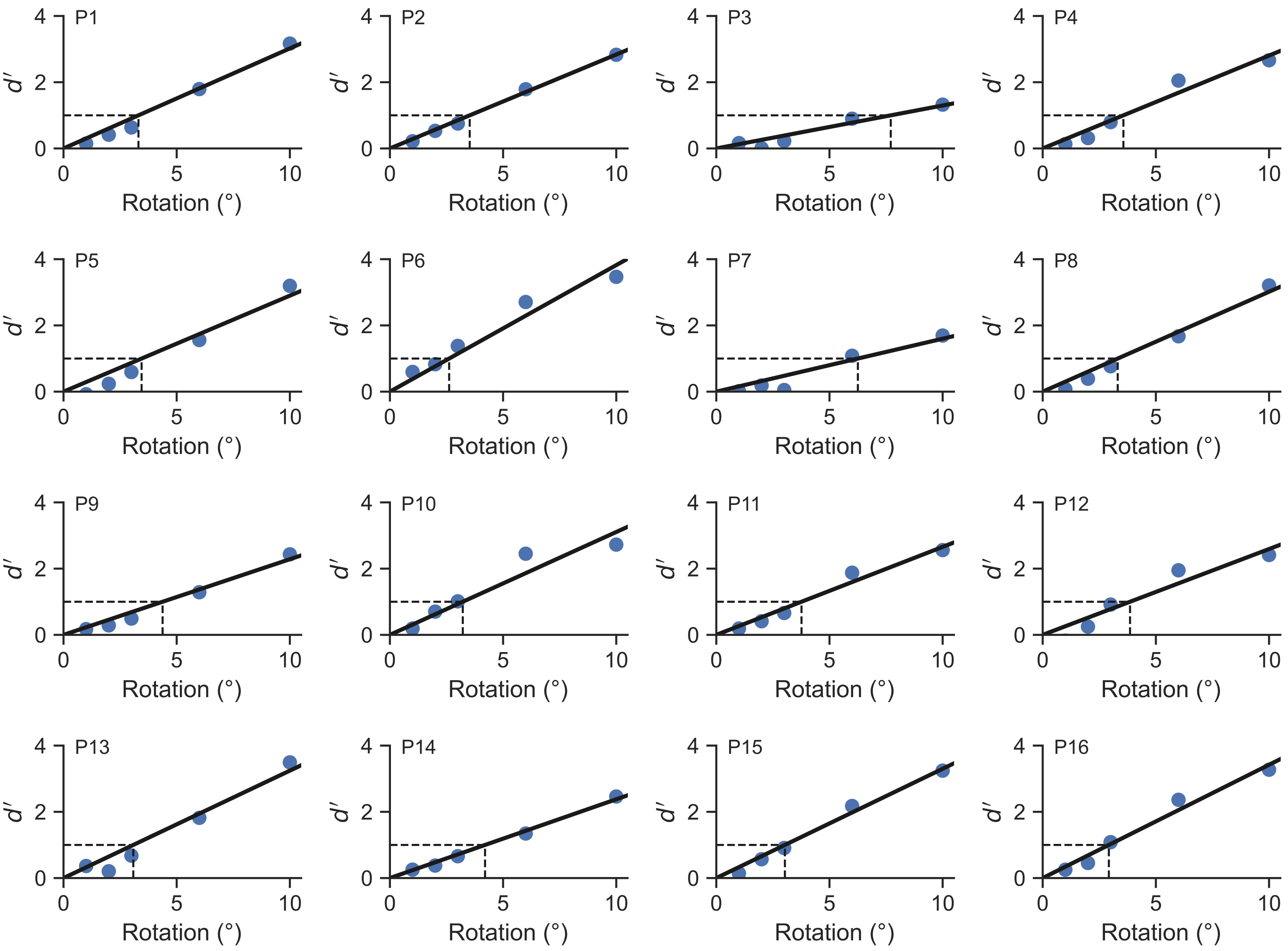
P1
P2
P3
P4
P5
P6
P7
P8
P9
P10
P11
P12
P13
P14
P15
P16
$d'$
Rotation (°)

### S4 Figure

# CO

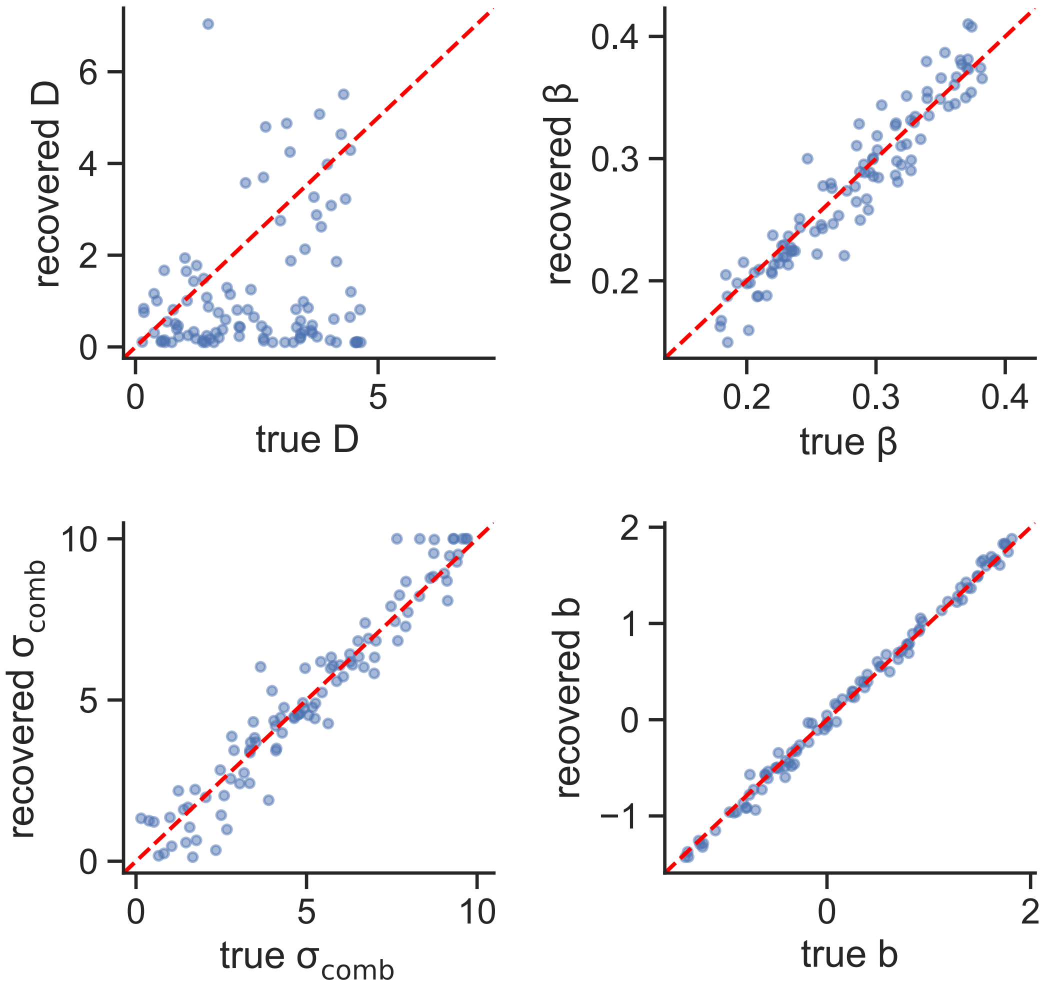

recovered D
true D
recovered β
true β
recovered $\sigma_{comb}$
true $\sigma_{comb}$
recovered b
true b

### S7 Figure

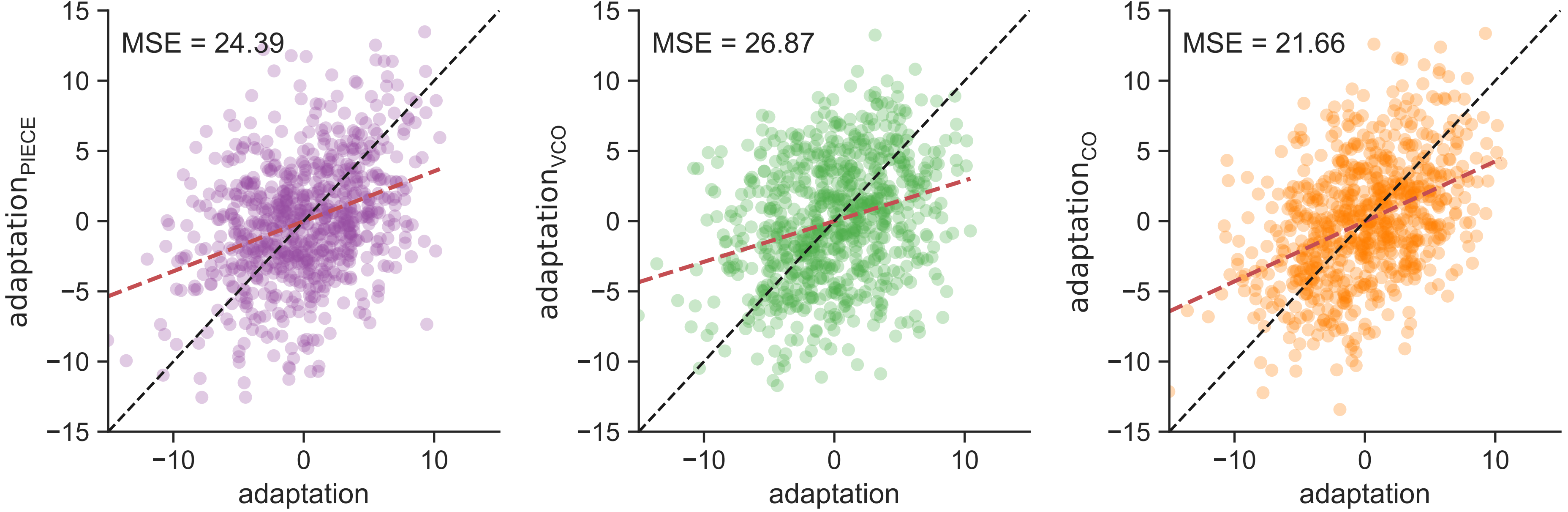

MSE = 24.39
MSE = 26.87
MSE = 21.66
adaptation_PIECE
adaptation_VCO
adaptation_CO
adaptation
adaptation
adaptation
15
10
5
0
−5
−10
−15
−10
0
10
