## Supplementary material for "Distinct computations underlie movement error parsing during implicit motor adaptation and perturbation detection": S3 Table

### Adaptation Task

| Model | Parameter | Mean [95% CI] |
|---|---|---|
| PIECE | $\alpha$ | 0.571 [0.466, 0.676] |
| | $\beta$ | 0.246 [0.225, 0.268] |
| | $\sigma_p$ | 0.195 [0.125, 0.266] |
| | $\sigma_{\text{pert}}$ | 100.731 [85.621, 115.842] |
| | $b$ | 0.167 [-0.360, 0.695] |
| VCO | $D$ | 5.890 [4.283, 7.557] |
| | $\beta$ | 0.209 [0.188, 0.231] |
| | $\sigma_v$ | 6.630 [4.762, 8.499] |
| | $b$ | 0.170 [-0.335, 0.675] |
| CO | $D$ | 1.651 [0.897, 2.406] |
| | $\beta$ | 0.301 [0.271, 0.331] |
| | $\sigma_{\text{comb}}$ | 3.899 [2.575, 5.223] |
| | $b$ | 0.176 [-0.327, 0.679] |

### Signal-Detection Task

| Model | Parameter | Mean [95% CI] |
|---|---|---|
| PIECE | $\alpha$ | 1.486 [1.074, 1.897] |
| | $\beta$ | 0.090 [0.225, 0.268] |
| | $\sigma_p$ | 3.289 [2.799, 3.779] |
| | $\sigma_{\text{pert}}$ | 4.540 [2.756, 6.324] |
| | $b$ | -0.263 [-0.790, 0.254] |
| VCO | $D$ | 3.964 [3.589, 4.340] |
| | $\sigma_v$ | 2.469 [2.137, 2.801] |
| | $b$ | -0.088 [-0.686, 0.509] |
| CO | $D$ | 3.475 [3.132, 3.817] |
| | $\sigma_{\text{comb}}$ | 2.881 [2.554, 3.222] |
