## Supplementary material for "Distinct computations underlie movement error parsing during implicit motor adaptation and perturbation detection": S5 Figure

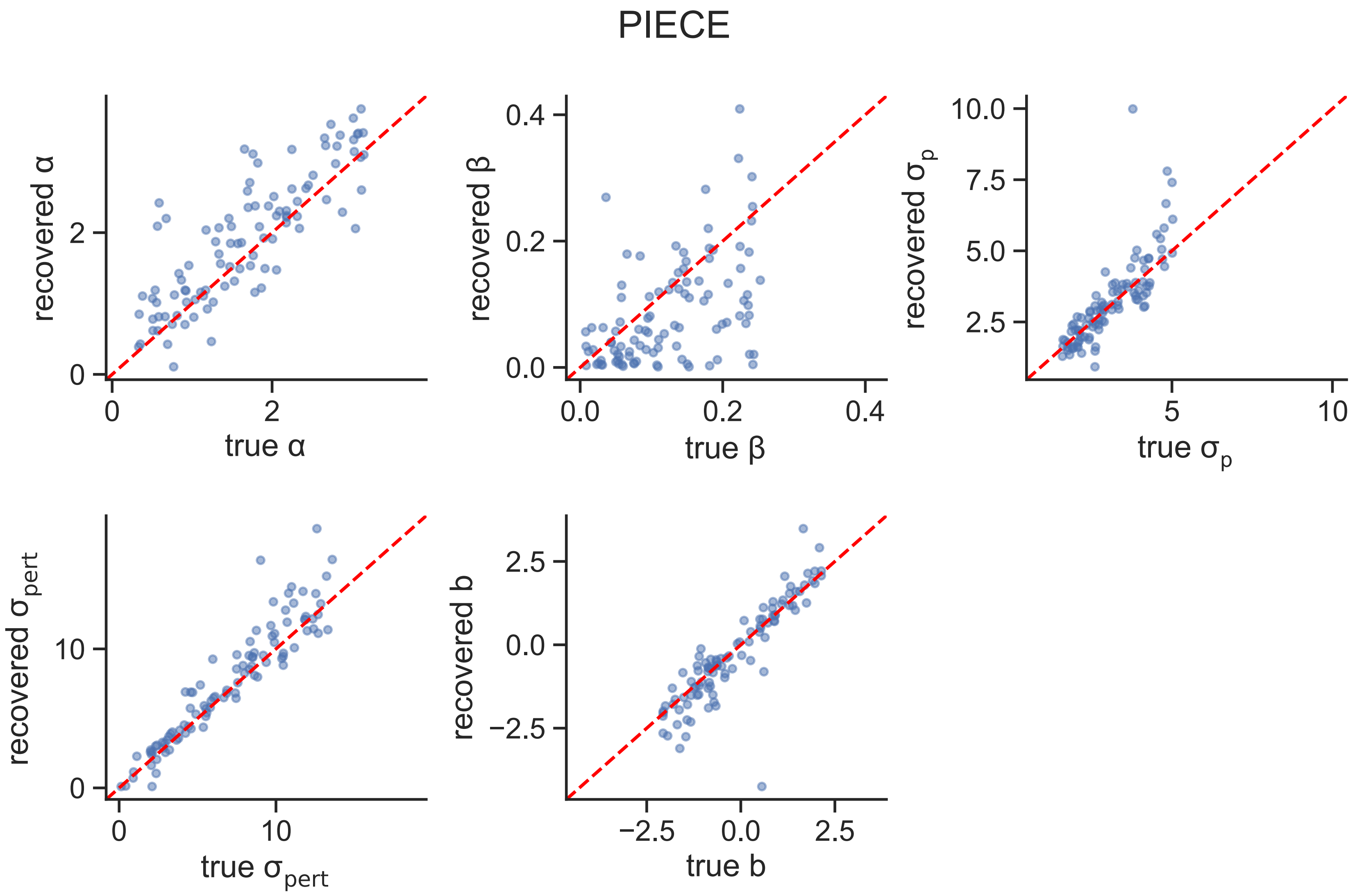

PIECE
recovered α
true α
recovered β
true β
recovered σp
true σp
recovered σpert
true σpert
recovered b
true b
