## Supplementary material for "Distinct computations underlie movement error parsing during implicit motor adaptation and perturbation detection": S6 Figure

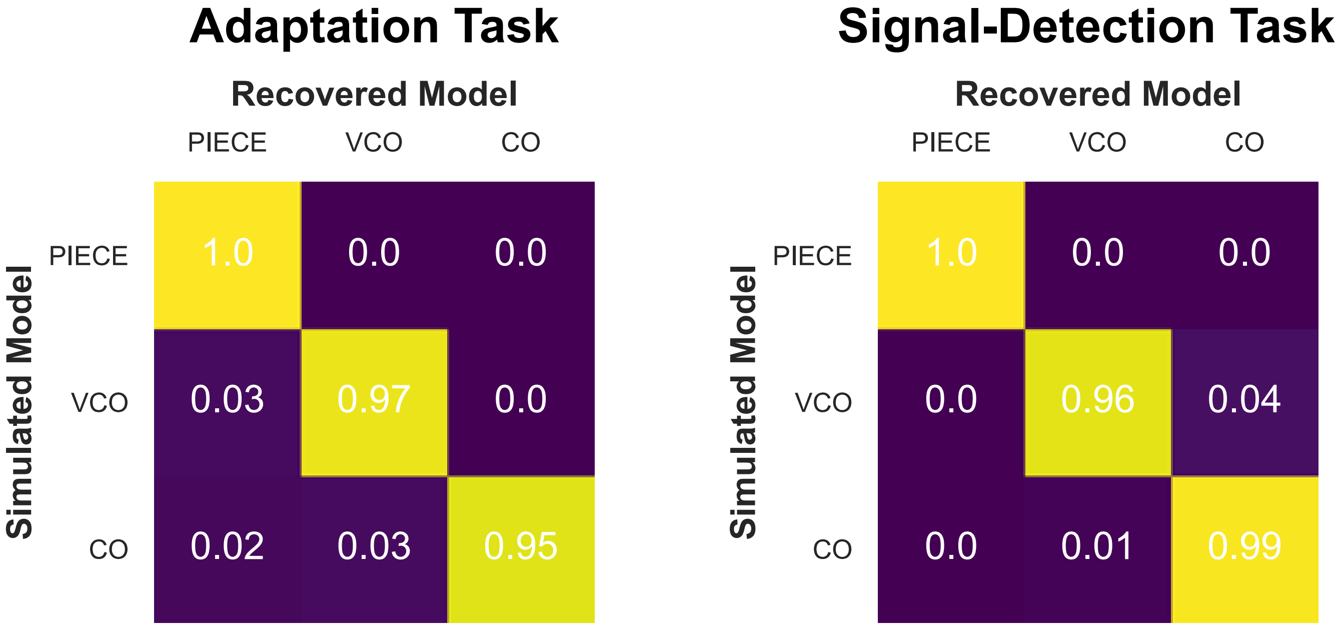
Adaptation Task
Recovered Model
PIECE
VCO
CO
Simulated Model
PIECE
VCO
CO
1.0
0.0
0.0
0.03
0.97
0.0
0.02
0.03
0.95
Signal-Detection Task
Recovered Model
PIECE
VCO
CO
Simulated Model
PIECE
VCO
CO
1.0
0.0
0.0
0.0
0.96
0.04
0.0
0.01
0.99
