## Supplementary material for "Distinct computations underlie movement error parsing during implicit motor adaptation and perturbation detection": S8 Figure

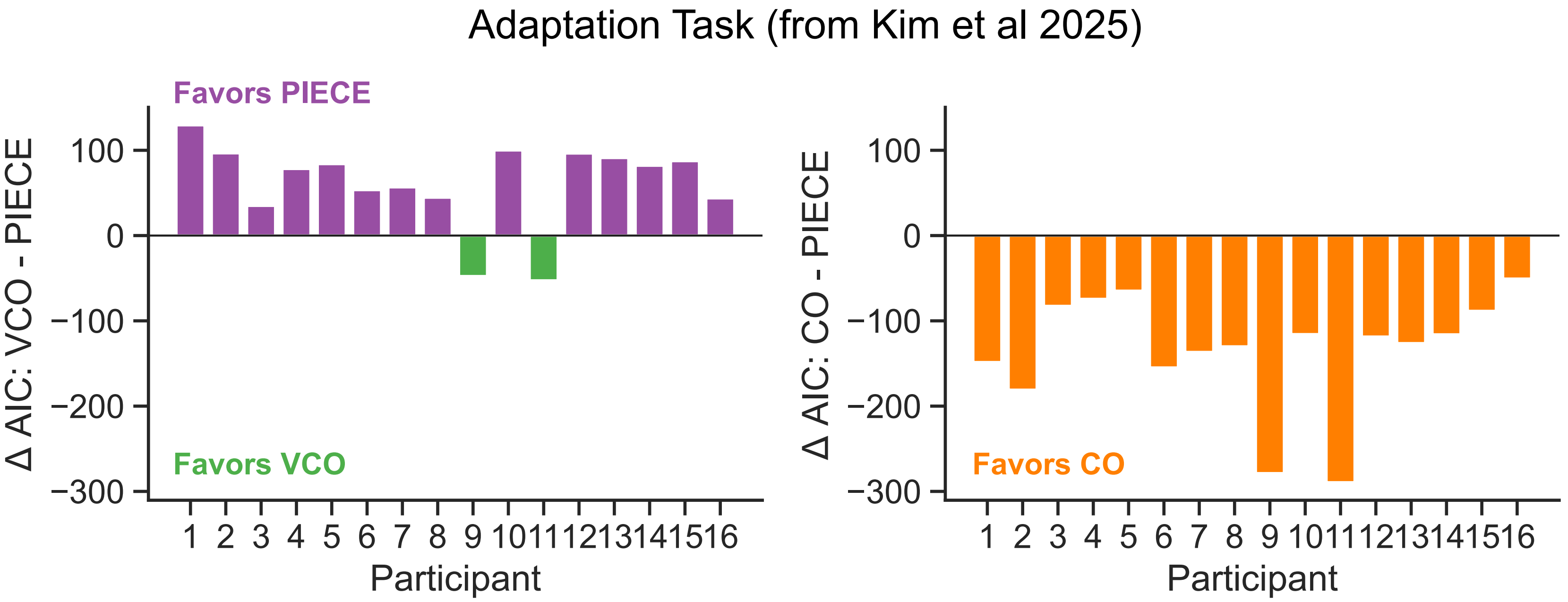
Adaptation Task (from Kim et al 2025)
Favors PIECE
Favors VCO
Δ AIC: VCO - PIECE
100
0
−100
−200
−300
1 2 3 4 5 6 7 8 9 10 11 12 13 14 15 16
Participant
Favors CO
Δ AIC: CO - PIECE
100
0
−100
−200
−300
1 2 3 4 5 6 7 8 9 10 11 12 13 14 15 16
Participant
